# iPSC-derived MSC secretome activates distinct neuroprotective pathways in a preclinical model of Parkinson’s disease

**DOI:** 10.1101/2025.11.07.687231

**Authors:** Ana Marote, Filipa Ferreira-Antunes, Sandra Barata-Antunes, Bárbara Mendes-Pinheiro, Carina Soares-Cunha, Ana Verónica Domingues, Stephanie Oliveira, Sandra Anjo, Daniela Monteiro, Patrícia Patrício, Bruno Manadas, Andreia Castro, Luísa Pinto, Belém Sampaio-Marques, António J Salgado

## Abstract

Parkinson’s disease (PD) is characterized by a progressive loss of dopaminergic neurons, which current therapies fail to address. Mesenchymal stem cells (MSCs) secretome holds promising therapeutic potential; however, the gold standard source - bone marrow (BM) - face scalability limitations. Induced pluripotent stem cells (iPSCs)-derived MSCs (iMSCs) are a rejuvenated and clinically attractive source, yet the effects of their secretome have not been studied in PD. Here, we directly compare the effects of BM-MSCs and iMSCs secretome in a 6-OHDA rat model. Despite few differences in secretome composition, both improved motor function and partially ameliorated anhedonia, but only iMSCs secretome significantly preserved dopaminergic neurons. Proteomic analysis revealed that BM-MSCs secretome activates antioxidant and proteostasis pathways, whereas iMSCs secretome activate broader pro-survival and regenerative pathways, including NRF2-mediated oxidative stress response and mTOR. In addition, iMSC-secretome treatment modulated glutamatergic and GABAergic neurotransmission, suggesting restoration of synaptic homeostasis beyond dopaminergic preservation. These findings demonstrate that iMSCs secretome exerts robust neuroprotective effects and shed light on the mechanisms underlying this scalable and clinically translatable therapy for PD.

## Introduction

Parkinson’s disease (PD) is a neurodegenerative disorder characterized by motor and non-motor symptoms that arise from neuropathological alterations in several neurotransmitter systems, including α-synuclein (asyn) accumulation and dopaminergic neuron loss in the nigrostriatal pathway [1]. Dopamine (DA) replacement therapy is initially effective in modulating motor symptoms, but long-term use is often associated with complications [2]. Therefore, the development of therapeutic strategies able to modify disease progression remains a major challenge [3].

Mesenchymal stem cells (MSCs) have shown promising therapeutic effects in several preclinical models of PD, mainly mediated by the neuroprotective and immunomodulatory activity of MSCs secreted factors [4–9]. In 6-hydroxydopamine (6- OHDA)-lesioned rodents, local administration of MSCs secretome improves motor function and reduces nigrostriatal tract cell loss[8, 9]. A similar neuroprotective effect is observed in an asyn overexpression rat model where MSCs secretome was able to reduce asyn mediated toxicity [5]. Recently, it was also shown that multiple systemic administrations of MSCs secretome in 6-OHDA-lesioned mice result in larger therapeutic effects, when compared to local delivery [8]. Targeted and non-targeted proteomic approaches in MSCs secretome have identified several neurotrophic factors that potentially mediate these neuropathological and behavioral improvements [10–12]. However, there is a lack of studies analyzing the target brain tissue, to determine how it receives and responds to these potential therapeutic factors.

MSCs used in previous PD studies have been mostly isolated from human bone-marrow (BM), which involves an invasive and painful procedure to the donor and presents decreased number, quality and proliferative capacity with donor’s age [13, 14]. Induced pluripotent stem cells (iPSCs) are a promising alternative source for obtaining large populations of MSCs (iMSCs), providing several advantages for clinical applications [15, 16]. First, iPSCs are an easily accessible and inexhaustible source of iMSCs [17]. Second, we and others have shown that iMSCs display a rejuvenated phenotype over BM-MSCs, which is translated into a superior proliferative capacity [10]. Finally, and importantly, iMSCs used in pre-clinical studies have yielded similar or even superior therapeutic activity in limb ischemia and bone regeneration [17]. A recent study has also demonstrated the therapeutic potential of iMSC transplantation in a PD mouse model [18]. Still, to our knowledge, the effects of iMSCs secretome on PD have not been addressed.

This study provides the first direct comparison of BM-MSCs and iMSCs secretomes regarding their composition and therapeutic efficacy in a unilateral 6-OHDA rat model of PD. We demonstrate, for the first time, that iMSCs secretome improves motor and non-motor function and reduces dopaminergic cell death in the *substantia nigra pars compacta* (SNc), with proteomic analysis suggesting distinct underlying molecular mechanisms between both secretomes.

## Results

### BM-MSCs and iMSCs cultured under serum-free conditions present differences in their secretome composition

Previous studies have shown that MSCs secretome composition and activity vary significantly depending on the source [19, 20]. To compare molecular profiles and identify potential therapeutically relevant factors, targeted and untargeted proteomic analyses were performed on triplicate BM-MSCs and iMSCs secretome samples.

MSCs secretome data set resultant from mass spectrometry analysis was first analyzed using STRING, which revealed a network comprising 86 nodes and 303 edges, with an average node degree of 7.05 and an average local clustering coefficient of 0.602, indicating a non-random, functionally interconnected network (Fig.1A). Further application of DBSCAN clustering method identified ten clusters based on the local node density. Fig.1A depicts five of these clusters, containing more than three proteins. Cluster 1 comprises 11 extracellular matrix (ECM) constituents (collagen subunits and fibronectin). ECM components influence neuronal survival and repair, and their presence in the network suggests that MSCs secretome may contribute to ECM remodeling, supporting neuroprotection and tissue repair [21]. Cluster 2 contains 15 proteins involved in protein folding (Calreticulin (CALR), Protein disulfide-isomerase A3 (PDIA3), Prolyl 4-hydroxylase subunit beta (P4HB), peptidyl-prolyl cis-trans isomerase B (PPIB), Heat Shock Protein 90 Beta Family Member 1 (HSP90B1), Heat shock protein 90 alpha family class B member 1 (HSP90AB1), heat shock protein family A (Hsp70) member 5 (HSPA5), Heat shock protein family A (Hsp70) member 8 (HSPA8)) and glycolytic process (glyceraldehyde-3-phosphate dehydrogenase (GAPDH), enolase 1 (ENO1), pyruvate kinase (PKM), triosephosphate isomerase (TPI1) and lactate dehydrogenase A (LDHA)). These proteins have important roles in proteostatic mechanisms, frequently altered in neurodegenerative disorders such as PD [22, 23]. Cluster 3 contains 12 proteins, mainly involved in translation (eukaryotic initiation factor 4A-I (EIF4A1), eukaryotic translation initiation factor 5A (EIF5A), eukaryotic elongation factor 2 (EEF2), ribosomal protein S23 (RPS23), ribosomal protein S9 (RPS9), ribosomal protein S18 (RPS18), large ribosomal subunit protein eL14 (RPL14), ribosomal protein S11 (RPS11), ubiquitin-ribosomal protein eL40 fusion protein (UBA52)) and structural constituents of chromatin (H4 clustered histone 6 (H4C6), histone H3.2 (H3C13), histone H2A type 2-A (H2AC18)), further suggesting that the majority of proteins identified can be a promising approach to target protein homeostasis dysregulation, associated with PD. Cluster 4 and 5 comprise proteins related to tubulin and actin cytoskeleton, respectively. These include tubulin subunits (tubulin beta-4B chain (TUBB4B), tubulin beta chain (TUBB), alpha-tubulin-1A (TUB1A), tubulin alpha-1C chain (TUBA1C), tubulin beta-6 chain (TUBB6), tubulin beta-2B chain (TUBB2B)) and proteins involved in actin cytoskeleton organization (cofilin-1 (CFL1), profilin-1 (PFN1), adenylyl cyclase-associated protein 1 (CAP1) and myosin-9 (MYH9)). Other proteins identified in this dataset display catalytic and antioxidant activity, namely peroxiredoxin-1 (PRDX1), thioredoxin (THIO), and glutathione S-transferase P (GSTP1), which is in line with previous reports on MSCs secretome, showing a strong composition in antioxidant molecules among different sources [19].

**Figure 1.**
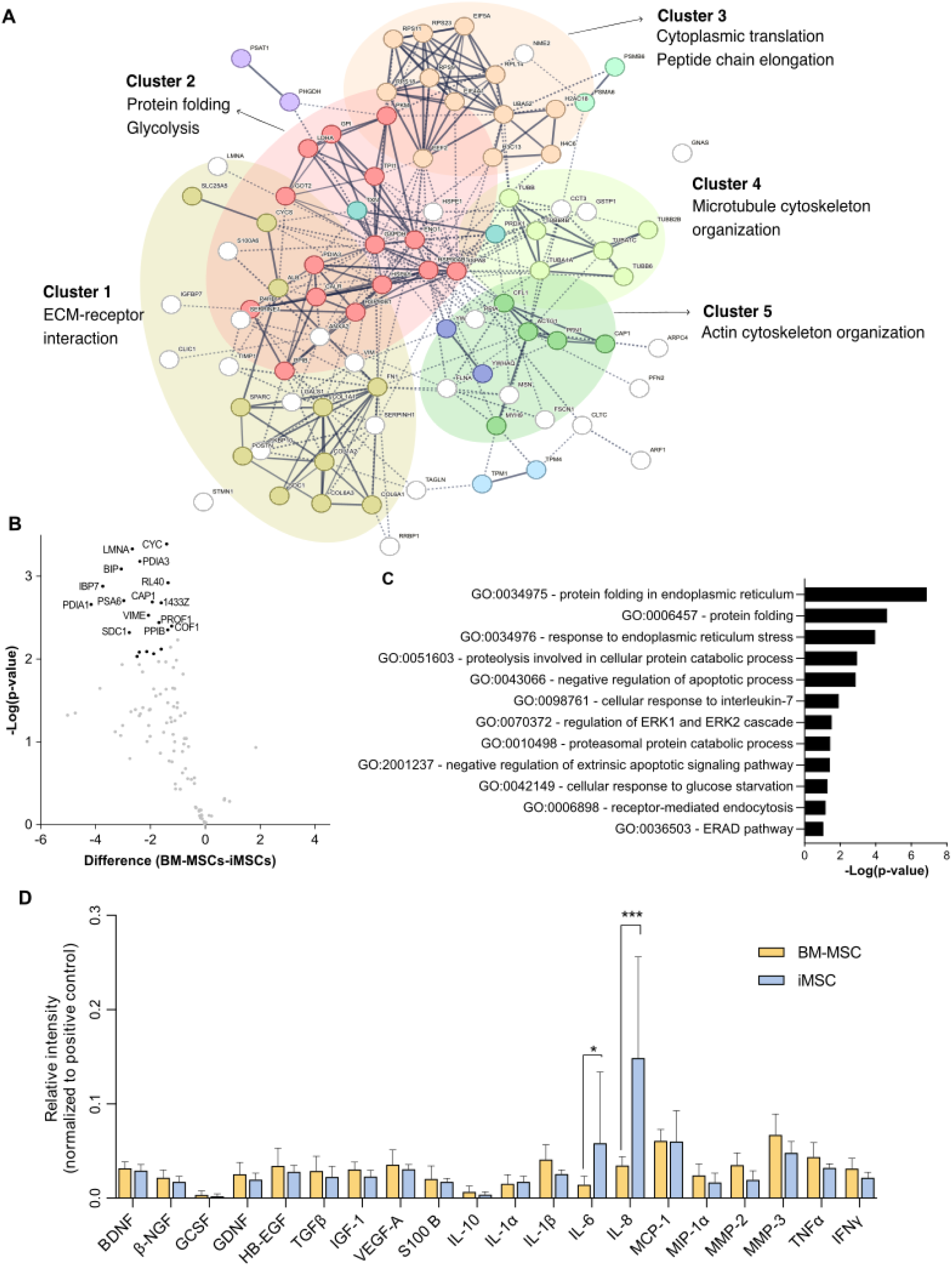
Comparative analysis of BM-MSC and iMSC secretome composition. **A.** STRING network analysis of proteins identified in BM-MSC and iMSC secretome by combined targeted and untargeted proteomic approaches (Average node degree of 7.05 and an average local clustering coefficient of 0.602). DBSCAN clustering method identified ten clusters based on the local node density and most enriched clusters are depicted in the figure. **B.** Volcano plot displaying differential protein abundance between BM-MSC and iMSC secretome. Proteins significantly enriched in iMSCs (FDR < 0.01) are expressed in the graph. **C.** Over-representation analysis of cellular component terms using ConsensusPathDB, highlighting enrichment in protein-folding- related components. **D.** Quantification of array spot intensity for each source after pooling the values from the three donors. Data is presented as mean ± SEM of relative intensity for each protein, normalized to positive control intensity. *p<0.05, ***p<0.001 for Sidak’s multiple comparisons test between BM-MSCs and iMSCs CM.

Regarding differences found between BM-MSCs and iMSCs secretome, only a limited number of proteins were significantly enriched in iMSCs secretome (FDR < 0.01; Fig.1B, Fig.S1A). Among those, are ER-associated proteins (endoplasmic reticulum chaperone BiP (BIP / HSPA5), protein disulfide-isomerase A1 (PDIA1), protein disulfide-isomerase A3 (PDIA3), PPIB, peptidyl-prolyl cis-trans isomerase FKBP10 (FKBP10)) proteosome subunits (proteasome subunit alpha type-6 (PSMA6), proteasome subunit beta type-6 (PSMB6)), and cytoskeletal or regulatory proteins (prelamin-A/C (LMNA), vimentin (VIM / VIME), syndecan-1 (SDC1), 14-3-3 protein zeta/delta (YWHAZ / 14-3-3ζ / 1433Z), insulin-like growth factor-binding protein 7 (IGFBP7 / IBP7)). Functional enrichment analysis through DAVID analysis (Fig. 1C), confirmed that these proteins participate in ER protein folding, response to ER stress, proteasomal catabolic process, and anti-apoptotic regulation.

To assess whether low-abundance cytokines and growth factors were also differentially secreted, a membrane-based antibody array was used (Fig.1D, Fig.S1B and S1C). Among the 20 proteins assessed, only IL-6 and IL-8 exhibited significant differences between sources, with iMSCs secretome showing higher levels of both (Fig.1D). IL-6 is a multifunctional cytokine secreted by neurons and glial cells, playing a vital role in neuronal development and differentiation [24, 25]. The secretion of key growth factors, including brain-derived neurotrophic factor (BDNF), glial cell line-derived neurotrophic factor (GDNF), insulin-like growth factor 1 (IGF-1), heparin-binding epidermal growth factor-like growth factor (HB-EGF) and growth factor beta (TGF-β), remained comparable between BM-MSCs and iMSCs. Similarly, other analyzed molecules such as macrophage inflammatory protein-1 alpha (MIP-1α), matrix metallopeptidase 2 (MMP-2), matrix metallopeptidase 3 (MMP-3,) S100 Calcium Binding Protein B (S100B), transforming Growth Factor alpha (TGFα), vascular endothelial growth factor A (VEGF-A), interferon gamma (IFNγ), interleukin 10 (IL-10), interleukin 1 alpha (IL- 1α) and interleukin 1 beta (IL-1β) showed no significant differences between the two sources (Fig.1D, Fig.S1B and S1C).

These findings suggest that while BM-MSCs and iMSCs share similar secretory profiles, iMSCs secretome exhibit higher expression of therapeutically relevant molecules for PD, further supporting their potential as a therapy for the disease.

### iMSCs secretome treatment improves motor and non-motor behavioral function in 6-OHDA lesioned rats

We next assessed the effects of iMSCs secretome in an *in vivo* model of PD, and compared it with BM-MSCs secretome, whose therapeutic potential has been previously reported [7–9]. A unilateral 6-OHDA rat model was established by stereotaxically injecting 8 µg of 6-OHDA into the right medial forebrain bundle (MFB), thereby selectively targeting and destroying dopaminergic neurons within the nigrostriatal pathway. This approach reproduces the dopaminergic degeneration characteristic of PD on one side of the brain, while the contralateral hemisphere serves as an internal control (Fig.2A and B).

**Figure 2.**
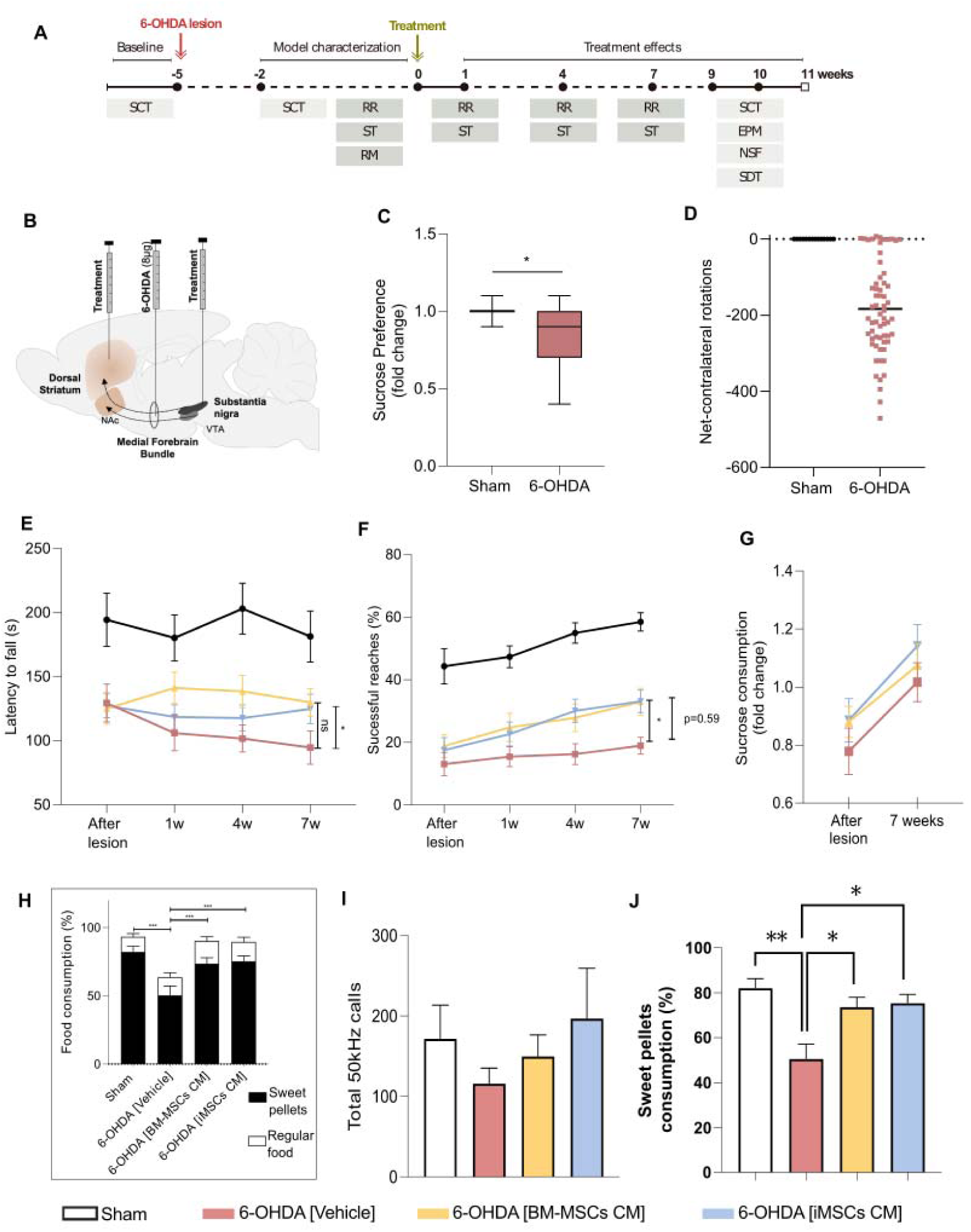
iMSC-derived secretome ameliorates motor and non-motor deficits in a 6-OHDA rat model of Parkinson’s disease. **A.** Experimental timeline showing motor (rotarod, staircase, apomorphine-induced rotations) and non-motor (sucrose consumption, SweetDrive, elevated plus maze, novelty-suppressed feeding) behavioral assessments following 6-OHDA lesion and secretome administration. **B.** Schematic of the unilateral 6-OHDA lesion model and injection sites for secretome delivery into the striatum and substantia nigra pars compacta (SNc). **C.** Sucrose consumption test showing fold change of preference after lesion relative to baseline (Mann–Whitney test, U = 138.5, *P* = 0.0312). **D.** Apomorphine-induced rotations confirm dopaminergic depletion compared to sham animals. **E.** Motor coordination and balance assessed by the rotarod test (latency to fall). Mixed-effects model (REML) with time, group, and their interaction as fixed effects revealed a significant main effect of group (F(3,50) = 5.588, *P* = 0.0022), but not time (F(2.500,111.7) = 1.767, *P* = 0.167) or interaction (F(9,134) = 1.369, *P* = 0.208). Post hoc Tukey correction. **F.** Fine motor performance in the staircase test (% successful reaches). Mixed-effects model showed significant main effects of time (F(1.929,86.81) = 28.57, *P* < 0.0001, η² = 0.060) and group (F(3,45) = 18.99, *P* < 0.0001, η² = 0.467), but no interaction (F(9,135) = 1.196, *P* = 0.303, η² = 0.008); Geisser–Greenhouse correction (ε = 0.643). Post hoc Tukey correction. **G.** Partial recovery of sucrose preference following secretome treatment. Mixed-effects model with time and group as fixed effects and subject as a random effect revealed a significant main effect of time (F(1,89) = 17.49, *P* < 0.0001), but not treatment (F(3,89) = 1.958, *P* = 0.126). Post hoc Tukey correction. **H.** Sweet Drive test (total food consumption). Two-way ANOVA revealed significant main effects of treatment (F(1,108) = 371.8, *P* < 0.0001), time (F(3,108) = 5.359, *P* = 0.0018), and their interaction (F(3,108) = 5.543, *P* = 0.0014). **i.** 50 kHz ultrasonic vocalizations showed no significant group differences (Kruskal– Wallis = 0.3973, *P* = 0.941; n = 4 groups, 51 total values). **J.** Sweet Drive test (sweet pellet preference). Group differences assessed using Brown–Forsythe and Welch’s ANOVA (Brown–Forsythe F(3,43.29)LJ=LJ7.420, *P*LJ=LJ0.0004; Welch W(3,29.27)LJ=LJ5.096, *P*LJ=LJ0.0059). Post hoc Dunnett’s T3 correction. *p<0.05 and **p<0.01. Data are shown as meanLJ±LJSEM (nLJ=LJ19–21 per group).

Three weeks after 6-OHDA lesion, animals underwent baseline assessment of motor and nonmotor functions (Fig.2A). Despite an initial weight loss (<10%), animals recovered well from the surgery and lesion (Fig.S2). In the sucrose consumption test, 6- OHDA lesioned animals displayed a reduced sucrose preference compared with non- lesioned animals (Fig.2C), consistent with a depressive-like phenotype previously described in this model [24]. Motor coordination and balance were evaluated using the rotarod test, where 6-OHDA lesion significantly reduced the latency to fall from the rotating beam (Fig.2E; p<0.05). In the staircase test, which reflects fine motor control in skilled reaching and forelimb dexterity, 6-OHDA lesion reduced animals’ ability to reach and eat the pellets placed on the staircase apparatus (Fig.2F). Apomorphine- induced rotational test (Rotameter) was used to assess dopaminergic imbalance and thus estimate lesion severity [25]. Of the 62 animals that received 6-OHDA, 50 presented intense turning behavior after apomorphine administration, as revealed by the increased number of contralateral rotations (Fig.2D), indicating an extensive dopaminergic denervation [25]. Together, these results confirm that the 6-OHDA model induced severe motor impairment and mild depressive-like behavior.

Animals were then divided into three treatment groups: vehicle (n=19, injected with Neurobasal A medium), BM-MSC (n=21, injected with BM-MSC secretome) and iMSC (n=21, injected with iMSC secretome). Although 12 animals did not exhibit rotational behavior, they were kept in the study since dopaminergic denervation can only be confirmed by histological analysis. Treatments were administered at the striatum and SNc of lesioned animals, and their effects were evaluated throughout the subsequent 11 weeks (Fig.2A). Secretome administration did not affect overall animal welfare, as body weight progressively increased in all groups (Fig.S2).

Both iMSCs and BM-MSCs secretome-treated animals exhibited improved performance in the Rotarod test compared with non-treated animals, although without statistical significance (Fig.2E). This improvement emerged one week after treatment administration and was sustained for up to seven weeks (Fig.2E). In the staircase test, treatment with iMSCs or BM-MSCs secretome significantly increased the successful reaches (p<0.05), similarly between both secretome groups (Fig.2F). These improvements were visible from the beginning of the post-treatment assessment, highlighting the neuroprotective potential of both iMSCs and BM-MSCs secretome.

While motor deficits are consistently reported in the unilateral 6-OHDA MFB-lesioned model, nonmotor alterations are less frequently examined and remain inconsistent, most likely due to protocol variability [26]. In this study, the reduced sucrose preference evidenced after 6-OHDA lesion, we observed a tendency that indicates possible reversion after BM-MSCs and iMSCs secretome-treatment (Fig.2G). A similar trend was observed in the sweet drive test, which comprises a multi-parametric analysis of anhedonia that integrates measurements of food preference in a non-aversive environment with recordings of 50 kHz ultrasonic vocalizations (USVs) associated with positive and pleasurable experiences [27, 28]. In comparison to non-lesioned animals, 6-OHDA-lesioned animals presented a decreased consumption of sweet pellets over regular food pellets (p<0.01) and a lower percentage of overall food consumption (Fig.2H; p<0.001), correlated with a reduction in 50kHz vocalizations (Fig.2I). Treatment administration, particularly iMSCs secretome partially reverted this phenotype, increasing both sweet pellet consumption (p<0.05) and total food consumption percentage (p<0.001) compared to non-treated 6-OHDA animals. Moreover, both MSCs and BM-MSCs secretome-treated animals showed a significant increase in sweet pellet consumption (Fig.2J; p<0.05) and overall food consumption (Fig.2H; p<0.001). Anxious-like behavior was also addressed using the Novelty Suppressed Feeding (NSF) test and Elevated Plus Maze (EPM) test, revealing no differences between lesioned and non-lesioned animals and administration of BM- MSCs or iMSCs secretome (Fig.S3A and S3B).

Overall, these results demonstrate that iMSC-derived secretome improves motor function in 6-OHDA-lesioned animals to a degree comparable with BM-MSC-derived secretome, supporting its use as a viable source for secretome with neuroprotective and regenerative potential. Moreover, these findings indicate that 6-OHDA lesion induces anhedonia, which is partially reversed by secretome treatment, highlighting the need to further explore the impact of MSCs secretome on non-motor symptoms of PD.

### iMSCs secretome protects dopaminergic neurons in the SNc, but has limited impact on striatal dopamine levels

At the end of the *in vivo* experiment, animals were allocated to one of two tissue processing protocols: whole brain fixation for subsequent sectioning and immunohistochemistry or macrodissection of specific brain regions for neurochemical profiling. Immunohistochemical quantification of tyrosine hydroxylase (TH) staining revealed a marked decrease in the density of fibers (>70%) and cell number (>90%) on the striatum and SNc, respectively, on 6-OHDA lesioned animals compared to sham animals (Fig.3C and 3G). Treatment with either BM-MSCs secretome or iMSC secretome promoted an increase on the TH+ fiber density in the striatum (Fig.3C), whereas only the administration of iMSCs secretome significantly prevented cell loss in the SNc (p<0.05; Fig.3G).

**Figure 3.**
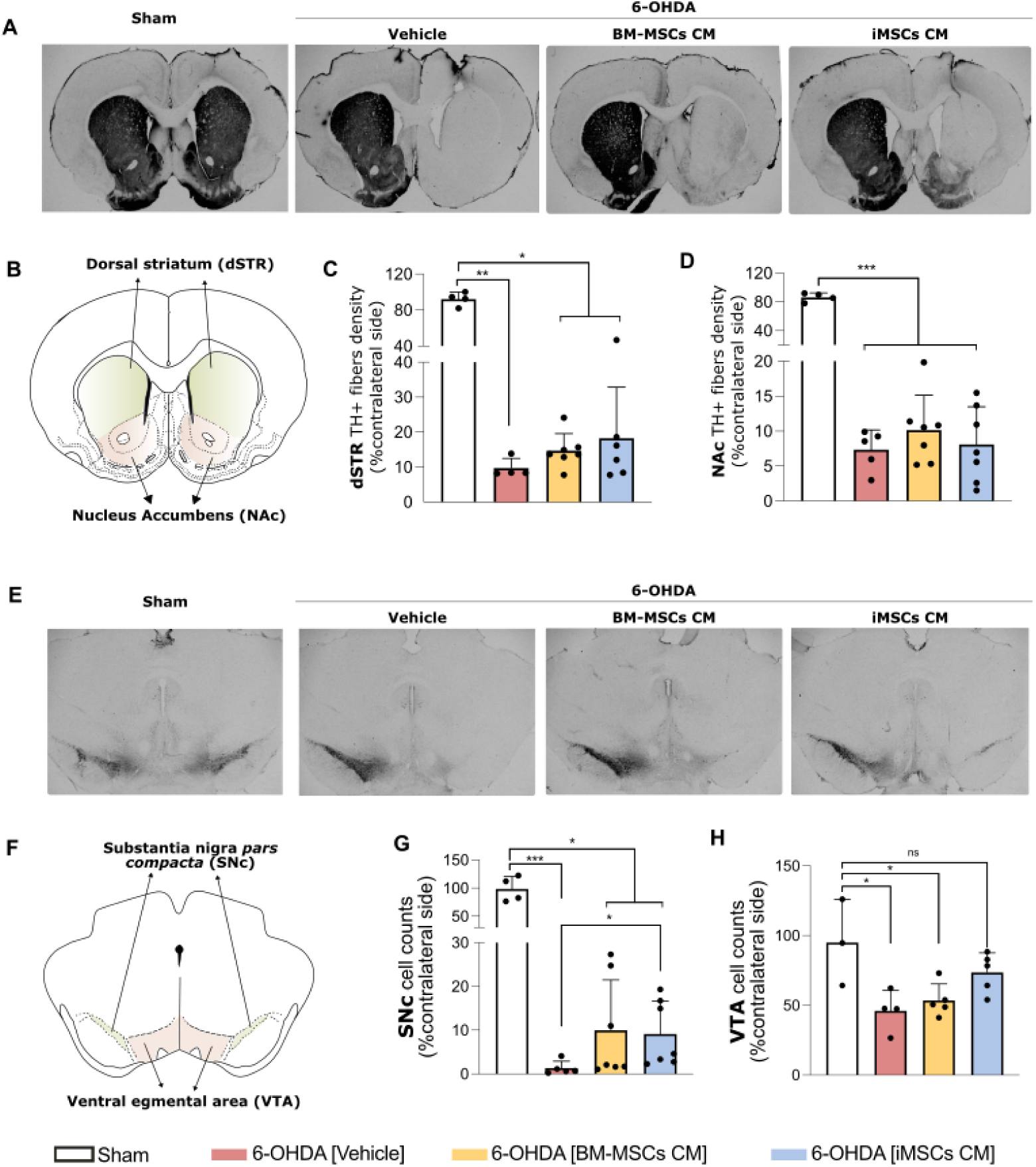
iMSC-derived secretome mitigates nigrostriatal and mesocorticolimbic dopaminergic neurodegeneration in 6-OHDA-lesioned rats. **A.** representative images of the tyrosine hydroxylase (TH) staining on the striatum; **B,** Diagram highlighting the dopaminergic projection in the striatum and nucleus accumbens (NAc). **C,** quantification of TH fibers density on the striatum, depicted as percentage over non-lesioned side analyzed with One-Way ANOVA (F(3,19) = 269.297, p<0.001, η² = 0.977). Post hoc comparisons were performed using Tukey’s correction**. D,** Quantification of TH+ fiber density in the NAc analyzed with One-Way ANOVA (F(3,17) = 79.328, p<0.001, η² = 0.933) and Welch’s ANOVA (W(3, 8.195) = 120.772, P < 0.001, η² = 0.933). Post hoc comparisons were performed using Dunn’s correction. **E,** Representative immunofluorescence images of TH staining in the Substancia nigra pars compacta (SNc) and Ventral tegmental area (VTA). **F.** Diagram highlighting dopaminergic populations in the SNc and VTA. **G.** Quantification of TH+ cells on the SNc, depicted as percentage over non-lesioned side analyzed with One-Way ANOVA (F(3,19) = 66.218, p<0.001, η² = 0.913). Post hoc comparisons were performed using Dunn’s correction**. H,** Quantification of TH+ cells in the VTA depicted as percentage over non-lesioned side analyzed with One-Way ANOVA (F(3,13) = 5.720, p=0.010, η² = 0.569). Post hoc comparisons were performed using Tukey’s correction.*p<0.05, **p<0.01 and ***p<0.001. Data are presented as meanLJ±LJSEM (nLJ=LJ6–9 animals per group).

Given the behavioral evidence of MSCs secretome effect in partially reversing anhedonia in 6-OHDA lesioned animals, we next examined the mesocorticolimbic pathway, which connects the ventral tegmental area (VTA) to the *nucleus accumbens* (NAc) in the ventral striatum and has fundamental roles in reward and aversion, usually impaired in depression associated to PD [29, 30]. Our results showed that 6-OHDA lesion significantly reduced the density of fibers in the NAc (>70%; Fig.3D; p<0.001) and reduced the number of cells in the VTA (>50%; Fig.3H; p<0.05) on 6-OHDA lesioned animals compared to sham animals. Treatment with iMSC secretome induced an increase on the TH+ cell counts in the VTA (Fig.3H), which might be correlated with the reversion of the anhedonic phenotype mentioned earlier.

Monoamines, including dopamine, norepinephrine and serotonin, are key neurotransmitters that regulate motor, cognitive, and emotional functions and have a critical role in PD associated motor and non-motor symptoms [31]. We used HPLC to quantify their striatal levels and thus evaluate the impact of the 6-OHDA lesion and subsequent treatments. As expected, 6-OHDA lesion induced a marked decrease in dopamine levels that was not reverted by administration of BM-MSCs or iMSCs secretome (Fig.4A; p<0.001). Interestingly, the analysis of dopamine turnover (DOPAC/dopamine ratio) in the contralateral hemisphere revealed a slight decrease in 6-OHDA lesioned animals, which was partially reverted by MSCs secretome treatment, even though without statistical significance. Regarding other monoamines, this quantification also revealed a significant decrease of norepinephrine levels on the striatum of lesioned animals either treated or non-treated (Fig.4C; p<0.001), whereas epinephrine and serotonin were not significantly altered.

**Figure 4.**
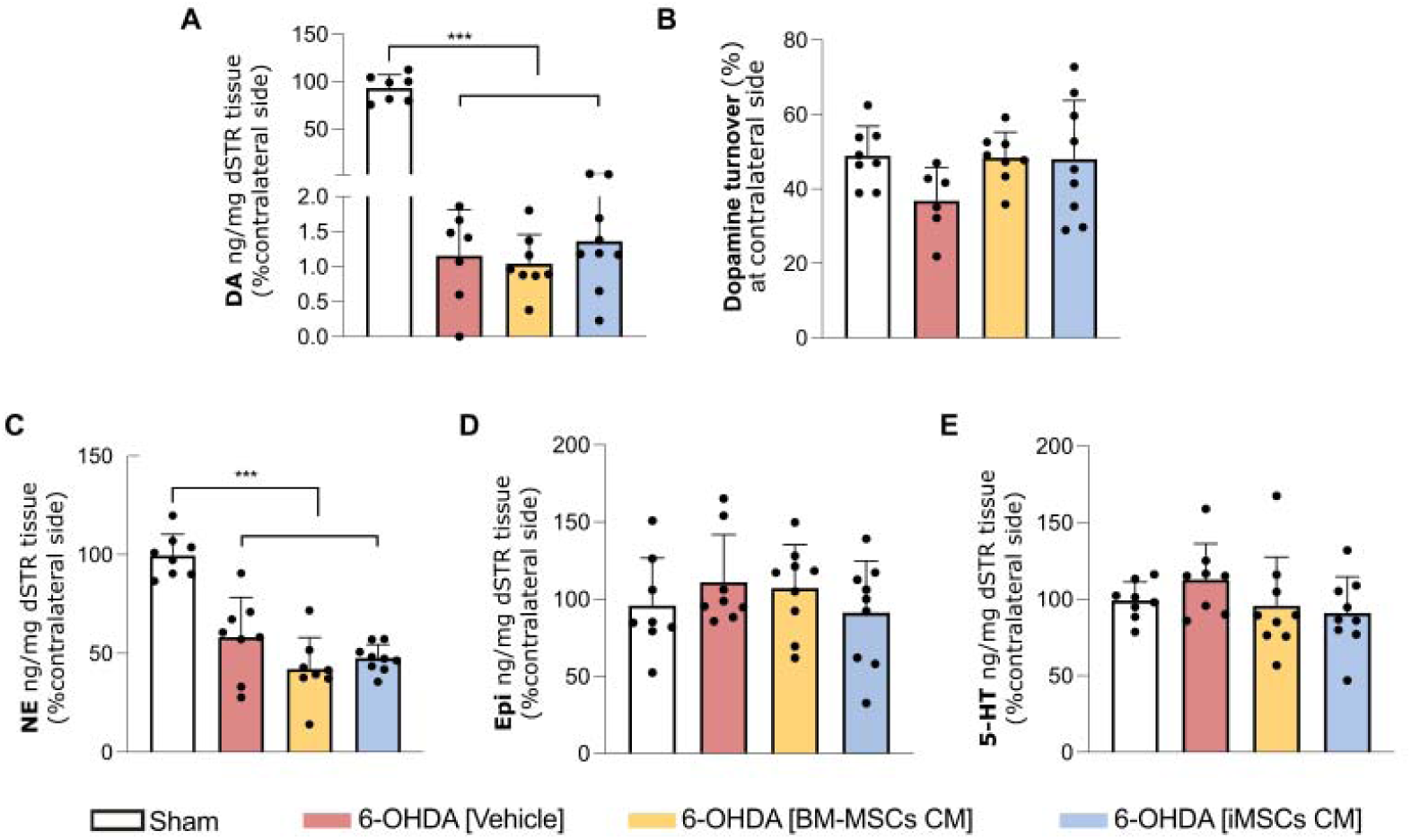
Neurochemical analysis of striatal catecholamine levels following 6-OHDA lesion and MSC-secretome treatment. **A** Quantification of dopamine (DA) release on the striatum performed by high-performance liquid chromatography, combined with electrochemical detection analyzed with One-Way ANOVA (F (3, 27) = 342,446, p<0.001, η² = 0.074). Post hoc comparisons were performed using Tukey’s correction. **B.** Dopamine turnover expressed as percentage over non-lesioned hemisphere analyzed with One-Way ANOVA (F (3, 27) = 1.858, p=0.161, η² = 0.171) and Welch’s ANOVA (W(3, 14.129) = 2.539, P = 0.098, η² = 0.171). **C.** Quantification of norepinephrine (NE) content in the striatum expressed as percentage over non-lesioned hemisphere analyzed with One-Way ANOVA (F (3, 29) = 26.966, p<0.001, η² = 0.736). NE content was also significantly reduced in all 6-OHDA-lesioned groups, regardless of treatment, compared to sham animals. Post hoc comparisons were performed using Tukey’s correction. **D**. Quantification of epinephrine (Epi) content in the striatum expressed as percentage over non-lesioned hemisphere analyzed with One-Way ANOVA (F (3, 30) = 0.773, p=0.518, η² = 0.072). **E.** Quantification of serotonin (5-HT) expressed as percentage over non-lesioned hemisphere analyzed with One-Way ANOVA (F (3, 30) = 1.272, p=0.302, η² = 0.113) and Welch’s ANOVA (W(3, 15.865) = 1.206, P = 0.340). Data are represented as meanLJ± SEM (n=7-8/group).

Together, histological and neurochemical analyses confirmed the severity of the nigrostriatal degeneration induced by 6-OHDA lesion. Importantly, the administration of iMSCs secretome partially attenuated neuronal loss in the SNc, consistent with the observed motor improvements. Moreover, iMSCs secretome also seems to have an effect in attenuating the degeneration elicited in the mesocorticolimbic pathway, specifically by preserving/increasing the number of dopaminergic cells in the VTA, which can be correlated with the slight reversion of the anhedonic phenotype in these animals.

### BM-MSCs and iMSCs secretome modulate distinct molecular pathways in the SNc of 6-OHDA lesioned rats

Until here, our results showed that both BM-MSCs and iMSCs secretome led to improvements at the behavioral and tissue level, even though iMSCs secretome induced more pronounced effects. To identify the underlying molecular mechanisms driving these differences, we conducted proteomic profiling of the SNc of a subset of animals included in the study.

Our proteomic analysis identified a total of 1877 proteins across SNc samples (n=3) from all groups. Subsequent hierarchical clustering of all quantified proteins (Fig.5A) revealed that secretome-treated biological replicates have some within-group variability, whereas Sham and non-treated groups display greater within-group consistency, as they cluster together. Importantly, MSCs secretome-treated samples were mostly separated from Sham and Vehicle groups along the primary dendrogram branches, suggesting that MSCs secretome induces proteomic remodeling. This was further confirmed by PCA, which revealed partial separation of experimental groups along PC1 (17.7% of variance) and PC2 (15.7% of variance) (Fig.5B). In this analysis, Sham and iMSCs secretome- treated samples clustered closely, suggesting proteomic similarity, whereas BM-MSC secretome-treated samples segregated distinctly.

**Figure 5.**
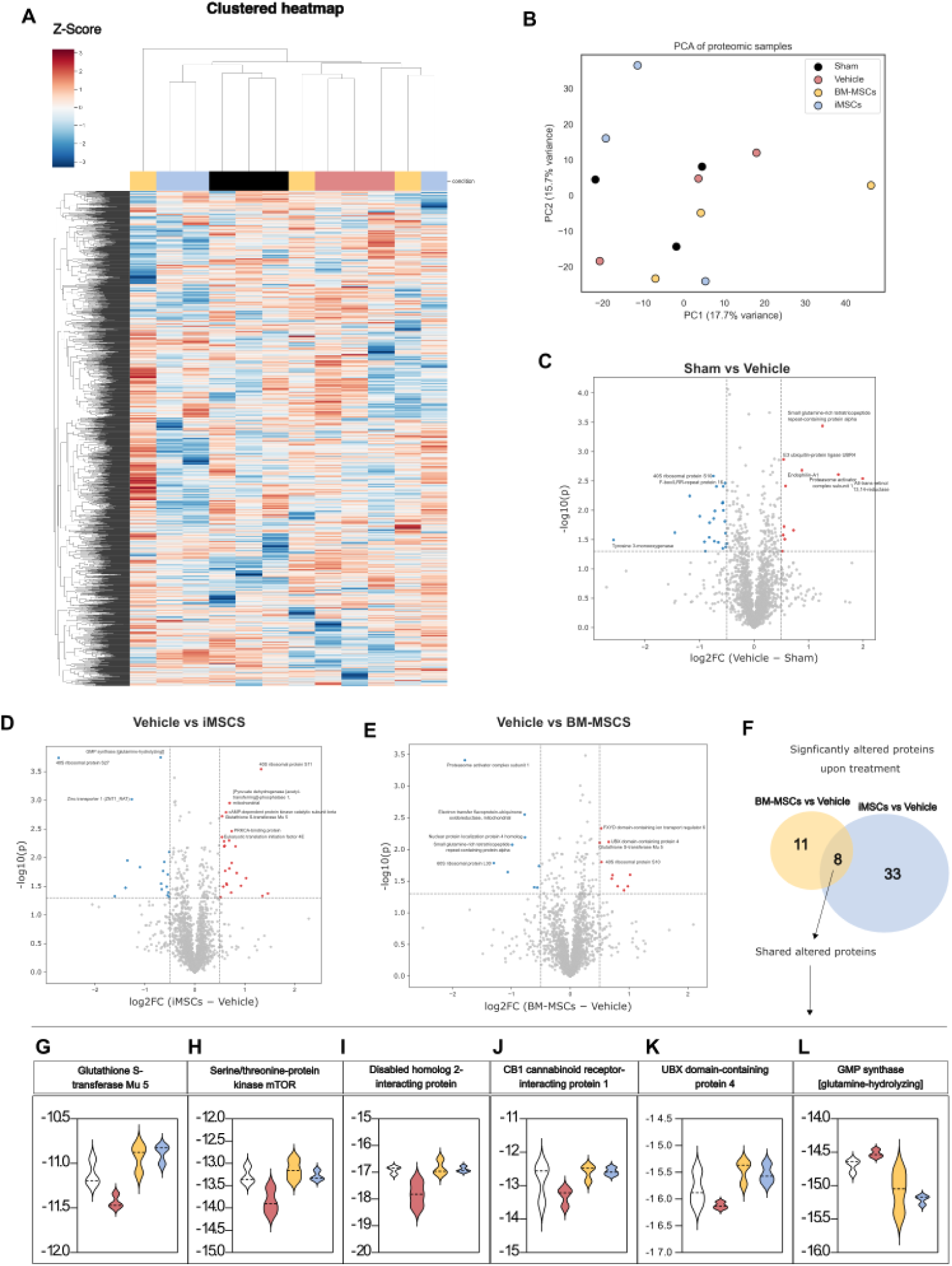
Proteomic profiling of the substantia nigra reveals distinct protein alterations induced by iMSC-secretome treatment. **A.** Hierarchical clustering of all quantified proteins; **B.** PCA, **C**. Volcano plot displaying differential protein abundance between Sham and Vehicle. Significantly altered proteins were defined as those with Benjamini–Hochberg FDR–adjusted p value < 0.05 in pairwise t-tests and an absolute log2 fold-change threshold of ≥ 0.5 (∼1.4-fold) and are represented in the graph. **D**. Volcano plot displaying differential protein abundance between Vehicle and iMSCs. Significantly altered proteins were defined as those with Benjamini–Hochberg FDR–adjusted p value < 0.05 in pairwise t-tests and an absolute log2 fold-change threshold of ≥ 0.5 (∼1.4-fold) and are represented in the graph. **E**. Volcano plot displaying differential protein abundance between Vehicle and BM-MSCs. Significantly altered proteins were defined as those with Benjamini–Hochberg FDR–adjusted p value < 0.05 in pairwise t-tests and an absolute log2 fold-change threshold of ≥ 0.5 (∼1.4-fold) and are represented in the graph. **F.** Venn diagram illustrating significant altered proteins upon treatment in iMSCs and BM-MSCs compared to vehicle. **G**. Log2 normalized values of Glutathione S-transferase Mu 5. **H**. Log2 normalized values of Serine/threonine-protein kinase mTOR. **I**. Log2 normalized values of Disabled homolog 2-interacting protein. **J**. Log2 normalized values of CB1 cannabinoid receptor- interacting protein 1. **K**. Log2 normalized values of UBX domain-containing protein 4. **L**. Log2 normalized values of GMP synthase.

To further compare these proteomic changes, volcano plots were generated to visualize protein-level differences between experimental groups (Fig.5C-E, Fig.S4). Significantly altered proteins were defined as those with Benjamini–Hochberg FDR–adjusted p value < 0.05 in pairwise t-tests and an absolute log2 fold-change threshold of ≥ 0.5 (∼1.4- fold). In the sham versus vehicle comparison (Fig.5C), 35 proteins were significantly altered. Among these is tyrosine 3-monooxygenase (the systematic name for tyrosine hydroxylase, TH), which is significantly downregulated in vehicle animals and confirms the detection of dopaminergic neurons in this dataset. Consistent with heatmap and PCA, sham and iMSCs secretome-treated animals show only 19 significantly altered proteins (Fig.S4A). In contrast, in treatment versus non-treatment comparison, iMSCs secretome induces major changes in SNc proteomic profile, with 42 proteins significantly altered in comparison with vehicle animals (Fig.5E). BM-MSCs secretome induces less alterations in lesioned animals, with only 19 proteins displaying significantly altered protein levels in comparison to vehicle animals (Fig.5D), and 33 in comparison to sham (Fig.S4B).

Even though direct comparison between treatment groups revealed 33 significantly altered proteins (Fig.S4C), we focused on the proteomic changes induced by each treatment relative to non-treated animals to better characterize the effects of MSCs secretome in this model, and to identify unique and shared molecular signatures upon treatment. A Venn diagram (Fig.5F) illustrates this comparison, showing that there is a shared alteration in eight proteins in BM-MSCs and iMSCs secretome. Log2 normalized values of six of these proteins are depicted in figures 5G-L. Most of them are upregulated in treatment groups, except for GMP synthase (Fig.5L). This protein is involved in GMP production, which has been demonstrated to increase intracellular calcium that precedes cell death [32]. On the other hand, upregulated proteins in MSCs secretome-treated animals are likely mediators of neuroprotection, including glutathione s-transferase mu 5 (Fig.5G), Serine/threonine-protein kinase mTOR (Fig.5H), Disabled homolog 2 interacting protein (Fig.5I), CB1 cannabinoid receptor-interacting protein 1 (Fig.5J), and UBX domain-containing protein (Fig.5K).

Next, we performed functional enrichment analysis using different bioinformatic tools to identify the main pathways being modulated upon MSCs secretome treatment (Fig.6 and Fig.S4). First, we used Ingenuity Pathway Analysis (IPA) to address the main canonical pathways being modulated upon secretome treatment. Figure 6A depicts a comparative analysis of the pathways modulated upon lesion (Sham vs Vehicle) and upon iMSC-secretome treatment (vehicle vs iMSCs), showing a clear effect of iMSCs secretome in reverting the alterations induced by 6-OHDA lesion. Most pathways are upregulated upon iMSCs secretome treatment, except for CLEAR (Coordinated Lysosomal Expression and Regulation) pathway, which regulates lysosomal biogenesis and function. Among the upregulated pathways are IL-8 and mTOR signaling, endocannabinoid developing neuron pathway, actin cytoskeleton signaling, synaptogenesis and epithelial junction signaling, which evidence the broad pathways that are directly or indirectly modulated by iMSCs secretome administration (Fig 6A, Fig.S4D). On the other hand, this comparative analysis was not possible to be performed in BM-MSCs secretome-treated animals, probably due to the few numbers of altered proteins in the analysis. Indeed, the most significantly enriched pathway is the protein ubiquitination pathway (Fig.S4E), driven by upregulation of multiple proteosome subunits. Alongside proteostasis, glutathione metabolism and glutathione redox reactions are also enriched, pointing toward a concerted cytoprotective response focused on reducing proteotoxic and oxidative stress [33–35], which is distinct from iMSCs secretome induced changes (Fig.S4D).

**Figure 6.**
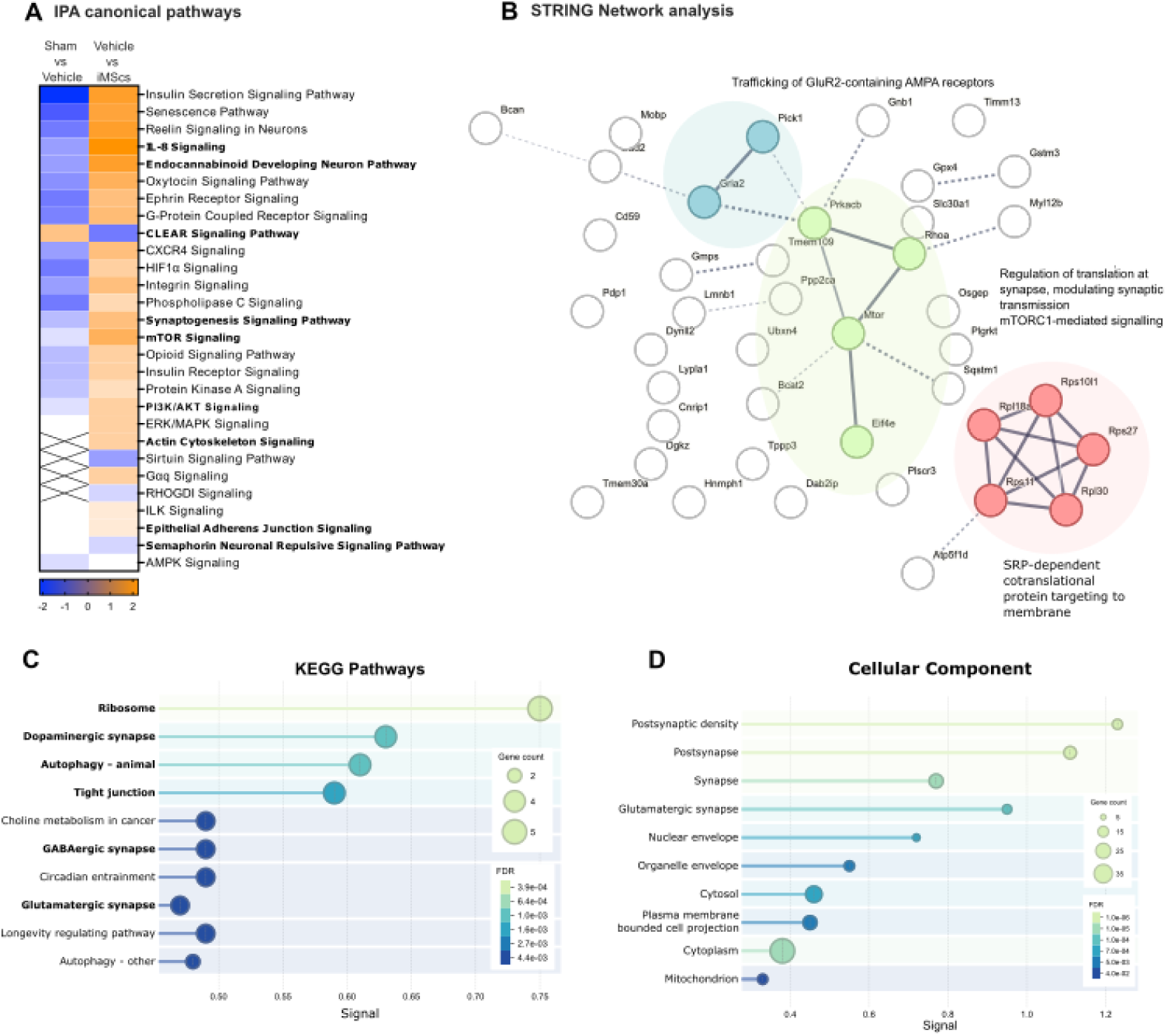
Comparative proteomic analysis of SNc tissue reveals distinct molecular pathways modulated by BM-MSCs and iMSCs secretome treatment. **A.** Ingenuity Pathway Analysis (IPA) to address the main canonical pathways being modulated comparing Sham vs Vehicle with Vehicle vs iMSC-secretome treatment. **B.** STRING network analysis of proteins identified in SNc of iMSC secretome treated rats. DBSCAN clustering method identified three clusters based on the local node density and most enriched clusters are depicted in the figure. **C.** KEGG Pathways analysis of SNc of iMSCs secretome treated animals. **D.** Cellular component analysis of SNc of iMSC secretome treated animals.

We proceeded this bioinformatic analysis looking only at the 42 proteins significantly altered by iMSC-secretome treatment. Using STRING network analysis, we identified three small clusters linked to Trafficking of GluR2-containing AMPA receptors, Regulation of translation at synapse, modulating synaptic transmission and SRP- dependent cotranslational protein targeting to membrane (Fig.6B). The identification of these clusters is in line with enriched KEGG pathways and cellular components, which evidence the involvement of these proteins in synaptic transmission, namely dopaminergic, GABAergic and Glutamatergic synapse (Fig.6C and D). For instance, there is an increased level of Glutamate decarboxylase 2 (Fig.S4F) and Glutamate receptor 2 (Fig.S4G). These findings indicate that the secretome exerts coordinated effects on synaptic structure and neurotransmission, not only enhancing dopaminergic signaling but also rebalancing excitatory (glutamatergic) and inhibitory (GABAergic) circuits [36, 37]. Moreover, the enrichment of autophagy-related proteins and ribosomal components (Fig.6C) further suggests an improvement in proteostasis and translational capacity, consistent with the observed regulation of mTOR and PKA signaling (Fig.6A).

Besides alterations at the neuronal level, iMSC secretome treatment is also likely to influence non-neuronal cell types. Indeed, alterations in tight junctions are evidenced in IPA (Fig.6A) and KEGG pathway (Fig.6C) analysis and have been reported in 6-OHDA animal models [38]. Finally, iMSCs secretome-treated animals also exhibited increased levels of Myelin-associated oligodendrocyte basic protein (MOBP) comparable to sham animals (Fig.S4H), suggesting a role in preserving oligodendrocyte integrity and myelin stability [39].

Together, these proteomic insights provide a mechanistic foundation for the observed histological and phenotypic improvements and highlight distinct therapeutic potential for BM-MSCs versus iMSCs secretome in the context of PD-related neurodegeneration.

## Discussion

The multifactorial etiology of PD requires a complex therapy that can address several dysregulated pathways and thus provide a reversion of the degenerative processes. The use of MSCs has long been studied as a therapeutic approach for PD [5, 8, 9, 17, 40], given their promising results in modulating several mechanisms associated with neurodegeneration, including oxidative stress, proteostasis and inflammation [17]. The possibility of obtaining iMSCs overcomes the isolation and culture hurdles of tissue- derived MSCs, such as BM-MSCs [10]. However, the differences regarding the therapeutic effects of iMSCs and BM-MSCs remain poorly understood, particularly in the context of PD. In this study, we sought to address the effects of iMSCs secretome in a PD rat model and provide a direct comparison with the gold-standard cell source, BM- MSCs regarding their composition and therapeutic potential.

We and others have shown that iMSCs display a rejuvenated phenotype in comparison to BM-MSCs, which is translated into a superior proliferative capacity before the induction of replicative senescence [10, 41]. In our previous study, iMSCs and BM- MSCs were expanded in hPL-supplemented culture media, which did not result in major alterations in their secretome profile [10]. Herein, iMSCs and BM-MSCs were expanded in a commercially available serum-free culture medium (Mesencult^TM^), which introduces less batch-to-batch variability and potential contamination of MSCs secretome with platelet-derived proteins [42]. In contrast to our previous study, there was a significant enrichment of 20 of the 86 proteins identified in proteomic analysis and increased IL-6 and IL-8 levels quantified in membrane antibody array in iMSCs secretome. Even though this was not the major focus of the present study, these results highlight the influence of MSCs expansion medium on secretome composition and the need to standardize and optimize protocols of MSCs expansion [43, 44].

Despite these differences in composition, both iMSCs and BM-MSCs secretome promoted significant improvements in motor performance in a 6-OHDA rat model of PD, evidencing, for the first time, the therapeutic potential of iMSCs secretome for PD. These results are in line with a recent study demonstrating the therapeutic potential of iMSCs systemic administration in a 1-methyl-4-phenyl-1,2,3,6-tetrahydropyridine (MPTP) mouse model [18]. While MSCs transplantation has been initially put forward as a therapeutic strategy for PD, it has been shown that MSCs beneficial effects are largely mediated by their paracrine signaling [7–9, 45]. The use of secretome as a cell transplantation-free approach represents a safer and more standardized alternative to MSCs transplantation, with greater potential for clinical translation [46].

Besides motor performance, our study further addressed the increasingly recognized non-motor component of PD in the *in vivo* model, demonstrating the effect of iMSCs secretome in partially reverting the anhedonic behavior of 6-OHDA lesioned animals. Accordingly, dopaminergic neurons in the VTA were preserved in iMSCs secretome- treated animals, which underscores the neuromodulatory potential of iMSC-secretome beyond the classic nigrostriatal pathway.

At the tissue level, only iMSCs secretome-treated animals exhibited a statistically significant preservation of dopaminergic neurons within the SNc. This finding contrasts with previous reports in which BM-MSCs secretome treatment demonstrated robust neuroprotection of dopaminergic neurons [8, 9]. One possible explanation lies in the specificities of BM-MSCs culture conditions used in this study, particularly MSC expansion media, which may have altered secretome profile. Alternatively, this result may also arise from the duration of the *in vivo* experiment. Here, histological analysis was performed 11 weeks post 6-OHDA lesion (versus 9 weeks in previous studies), a time-window during which BM-MSCs secretome may have not be sufficient to counteract the progressive neurodegenerative process.

A key innovative approach of this work lies in the proteomic characterization of the SNc, which provided a mechanistic insight of MSC secretome effects in this PD model. This analysis revealed distinct patterns of pathway modulation between treatment groups, underscoring the functional significance of the differences in secretome protein abundance between BM-MSCs and iMSCs. Importantly, before dissecting the potential mechanisms linked to each secretome source, it is critical to highlight the proteins that were modulated in both treatments, which may put forward shared effectors underlying MSC secretome effects in PD. Among the 19 proteins significantly altered in BM-MSCs secretome treated animals, 8 overlapped with those modulated by iMSCs secretome group (42 proteins in total). Notably, these alterations were observed when comparing treated versus non-treated animals, and treatment restored protein levels to those of sham animals, suggesting a therapeutic effect of these proteins in response to 6-OHDA insults.

Among the 8 proteins identified in both sources is Glutathione S-transferase Mu 5 (GSTM5), which participates in cellular defense mechanism against oxidative stress [47]. Reduced glutathione levels and impaired detoxification capacity are described as main drivers of SNc degeneration in PD [48, 49]. Accordingly, in our analysis, GSTM5 is decreased in response to 6-OHDA lesion (vehicle vs sham) and treatment with either BM-MSCs secretome or iMSC secretome-treated animals seems to actively potentiate this endogenous antioxidant mechanism. In line with this result, IPA analysis revealed an enrichment of Glutathione redox reactions and Glutathione-mediated detoxification in BM-MSCs secretome-treated animals, whereas iMSC-secretome treated animals display an enrichment of NRF2-mediated oxidative stress response. NRF2 activation upregulates antioxidant enzymes, including GSTM5, acting in cellular redox buffering, mitochondrial function and modulating inflammation, which are mechanisms that counteract the oxidative insults produced by dopamine metabolism in patients and by 6- OHDA lesion in animals [50, 51]. These findings are in line with a growing body of evidence showing that MSC secretome exert protective effects against oxidative stress, thus supporting neuronal survival after 6-OHDA insult [8, 9, 52].

Another key protein altered in both treatment groups is Serine/threonine-protein kinase mTOR, which is decreased in vehicle group. Indeed, overexpression of mTOR was shown to protect neurons from 6-OHDA toxicity, while inhibition of mTOR exacerbated dopaminergic neuronal death [53]. However, the role of mTOR modulation in PD remains controversial. While some authors suggest that upregulation of mTOR can suppress autophagy, thereby impair the clearance of misfolded proteins and contribute for disease progression, others suggest that mTOR inhibition can enhance autophagic removal of toxic aggregates [53, 54]. Besides this role in autophagy, mTOR activation has been shown to enhance intracellular signaling supporting cell survival, growth, and metabolic resilience in dopaminergic neurons [55–57]. Importantly, mTOR signaling pathway is also a highly enriched pathway in iMSCs secretome-treated animals, which can be advocate for the effect of iMSC secretome in preserving dopaminergic neurons after 6-OHDA lesion. In line with this increase in mTOR, autophagy was also evidenced in BM-MSCs secretome-treated animals (IPA) and in iMSC-secretome treated animals (KEGG pathway).

We also found upregulated CB1 receptor–interacting protein in secretome–treated animals, suggesting a role in endocannabinoid receptor regulation in the SNc [58, 59]. Once again, Endocannabinoid Developing Neuron Pathway was also enriched in iMSCs secretome-treated animals. The endocannabinoid system critically regulates basal ganglia function, synaptic plasticity, and neuroinflammation, processes disrupted in PD and following nigrostriatal injury [58, 60, 61]. In 6-OHDA models, CB1 receptor expression and endocannabinoid tone undergo lesion- and activity-dependent modulation [62, 63]. Functionally, CB1 signaling exerts context-specific effects: receptor antagonism can alleviate motor hyperinhibition, whereas CB1 agonists or endocannabinoid enhancers confer neuroprotection by dampening excitotoxicity, oxidative stress, and glial activation [58, 60]. The upregulation of a CB1-interacting protein in iMSC-treated animals may therefore reflect a compensatory mechanism that fine-tunes receptor trafficking and signaling, contributing to both neuromodulatory and neuroimmune balance within the substantia nigra.

In line with the differences in secretome composition and tissue-level effects in the PD model, our proteomic analysis further suggests iMSCs secretome treatment-specific alterations. For instance, there was an enrichment in IL-8 signaling pathway, consistent with the elevated IL-8 content observed in iMSC secretome. Besides its canonical role in inflammation, IL-8 has been described to mediate other effects that may be important in the context of PD, including neuroprotection, angiogenesis, and glial activity, which may contribute for the observed dopaminergic preservation [64, 65].

iMSCs secretome treatment may also elicit synaptic remodeling, supported by an upregulation of actin cytoskeleton signaling and downregulation of semaphorin neuronal repulsive signaling pathway, thereby favoring structural stabilization and synaptic connectivity. In addition, iMSCs secretome treatment appears to modulate glutamatergic and GABAergic neurotransmission. Dysregulation of excitatory/inhibitory balance has been investigated in PD and neurotoxin models, where dopamine depletion leads to excessive glutamatergic drive within basal ganglia circuits and secondary alterations in GABAergic tone [36]. These maladaptive changes contribute to abnormal motor output and excitotoxic stress contributing to a worsening of phenotype [36]. The observed modulation of proteins related to glutamate receptors, GABA receptor signaling and synaptic organization suggests that iMSC-secretome treatment may help restore synaptic equilibrium between excitatory and inhibitory systems by enhancing GABAergic transmission [66]. Thus, upregulation of glutamatergic and GABAergic transmission may be linked to the reversion of motor deficits even in the absence of measurable restoration of striatal dopamine levels which can also be justified by the sensitivity limits of neurochemical assays such as HPLC.

This proteomic snapshot provides a broad view of the molecular adaptations occurring after 6-OHDA injury and their modulation by the iMSC secretome. While it does not allow definition of a precise mechanistic pathway, it reveals coordinated regulation of pathways related to neuronal survival, oxidative stress, synaptic remodeling, and glial support. Indeed, several of the altered proteins are not exclusively neuronal, suggesting that secretome treatment may also modulate non-neuronal populations, including oligodendrocytes and other glial cells, which is aligned with previous studies [5, 67].

Future studies dissecting the contribution of individual secretome components or specific signaling nodes will be necessary to clarify causal relationships. However, accumulating evidence supports the notion that the therapeutic efficacy of MSC-derived secretome arises from the concerted action of its complex molecular constitution, rather than from isolated factors [68]. In this context, our data demonstrate that iMSC secretome exerts a broader and more effective modulation of neuroprotective and neurotransmission-related pathways than the BM-MSC secretome, indicating a superior neuromodulator potential and enhanced translational relevance for PD therapy. Moreover, the scalability, lower production cost, and reduced immunogenic and infectious risk associated with iMSC-derived secretome underscore its strong translational potential as a clinically viable, cell-free therapeutic strategy.

## Conclusion

This study provides compelling evidence that the therapeutic potential of MSC-derived secretome is strongly influenced by its cellular origin. While both BM-MSCs and iMSCs exhibit comparable secretory profiles, our data reveals that the iMSC secretome contains a higher abundance of key bioactive molecules involved in neuroprotection, oxidative stress mitigation, and neuromodulation. These compositional differences are reflected in the enhanced histological preservation of dopaminergic neurons and the more robust behavioral improvements observed in iMSC-treated animals. Moreover, pathway analysis suggests that the iMSC secretome exerts a broader impact on critical mechanisms underlying PD pathophysiology. In this regard, iMSCs represent a promising, scalable, and clinically relevant source of neuroprotective factors aimed at halting or reversing disease progression.

## Materials & methods

### MSCs culture and secretome collection

Human BM-MSCs (Lonza, Switzerland) from three donors were compared with iMSCs generated as previously described [10]. Both cell types were expanded in MesenCult™- ACF Plus Medium and passaged with Animal Component-Free Cell Dissociation Kit (STEMCELL Technologies, Canada). Secretome was collected at passages 8–9 after 72 h culture (5,000 cells/cm² for in vivo/protein array; 12,000 cells/cm² for proteomics), followed by 24 h conditioning in Neurobasal A medium with 1% kanamycin. Media were centrifuged (1,200 rpm, 10 min, 4 °C) to remove debris, concentrated 100× (5 kDa cut-off; Vivaspin) for in vivo/proteomics, and stored at –80 °C.

### Secretome proteomic analysis

For secreted protein preparation, an internal standard (IS; recombinant maltose-binding protein fused to green fluorescent protein, MBP-GFP) was added in equal amounts (1 µg) to each sample [69], and the supernatants with the secreted proteins were completely dried under vacuum using a Speedvac Concentrator Plus (Eppendorf). Supernatants were dried under vacuum (Speedvac Concentrator Plus, Eppendorf), pellets resuspended in SDS sample buffer and processed with ultrasonication (750 W ultrasonic processor) and 95 °C denaturation. Digestion was performed as described previously [70].

Protein quantification was performed by Sequential Windowed Acquisition of All Theoretical Mass Spectra (SWATH-MS). Samples were analyzed on a NanoLC™ 425 system coupled to a TripleTOF™ 6600 mass spectrometer (Sciex®) using: i) pooled samples analyzed by information-dependent acquisition (IDA), and ii) individual samples by SWATH-MS.

The DuoSpray™ ion source (25 µm ID hybrid PEEKsil/stainless steel emitter, ABSciex®) was used. Peptides were loaded onto a YMC-Triart C18 guard column (12 nm, S-3 µm, 5 × 0.5 mm) at 5 µL/min (5% mobile phase B, 4 min), then separated by micro-flow LC on a YMC-Triart C18 analytical column (150 × 0.3 mm) at 50 °C. The gradient was: 5–30% B (0–25 min), 30–90% B (25–26 min), 90% B (26–32 min), and 90–5% B (32–35 min), at 5 µL/min. Mobile phases: A = 5% DMSO, 0.1% formic acid in water; B = 5% DMSO, 0.1% formic acid in acetonitrile. Ion source settings: positive mode, 5500 V spray voltage, GS1 = 25 psi, CUR = 25 psi, source temperature = 100 °C. For IDA, MS scans (m/z 350–1250, 250 ms) were followed by 30 MS/MS scans (m/z 100–1500, 75 ms) per 2.8 s cycle. Ions (+2 to +5, ≥100 cps) were fragmented once before 15 s dynamic exclusion, using rolling collision energy with CES 5. For SWATH, 30 overlapping 25 Da windows covered m/z 350–1100. Each cycle began with a 50 ms survey scan (m/z 350–1250) followed by 50 ms MS/MS scans, total 2.8 s cycle time; collision energy was set for +2 ions with CES 15 [71].

Peptide identification and library generation were performed by searching all the IDA samples using the ProteinPilot™ software (v5.1, ABSciex®) with the following parameters: i) search against a database from SwissProt composed by Homo Sapiens (downloaded in September 2020) and MBP-GFP (IS) protein sequences; ii) acrylamide alkylated cysteines as fixed modification; and iii) trypsin as digestion type. An independent False Discovery Rate (FDR) analysis using the target-decoy approach provided with Protein Pilot software was used to assess the quality of the identifications and positive identifications were considered when identified proteins and peptides reached a 5% local FDR [72, 73].

Quantitative data processing was conducted using SWATH™ processing plug-in for PeakView™ (v2.0.01, ABSciex®) [74]. After retention time adjustment using the MBP- GFP peptides, up to 15 peptides, with up to five fragments each, were chosen per protein, and quantitation was attempted for all proteins in the library file that were identified from ProteinPilot™ search. Only proteins with at least one confidence peptide (FDR<0.01) in three biological replicates per condition and with at least three transitions were considered. Peak areas of the target fragment ions (transitions) of the retained peptides were extracted across the experiments using an extracted-ion chromatogram (XIC) window of 4 minutes with 100 ppm XIC width. The proteins’ levels were estimated by summing all the transitions from all the peptides for a given protein that met the criteria described above [75] and normalized to the levels of the IS of each replicate [69].

Protein-protein interaction network analysis was constructed using the online STRING database (https://string-db.org) version 12.0, depicting functional and physical protein associations with a high confidence level (0.7), organized into ten clusters, through DBSCAN clustering method (epsilon parameter = 3).

For the identification of differentially regulated proteins, the relative protein quantification was processed using the Perseus 1.6.14.0 Software (Max Planck Institute of Biochemistry). After data log2 normalization, Student’s t-test was used to test the differences between groups, and a false discovery rate (FDR or q-value) < 0.01 was considered significant. All proteomic data was displayed in a volcano plot according to their statistical p-value and their relative difference of abundance between both sources using Perseus 1.6.14.0 Software (Max Planck Institute of Biochemistry). The Database for Annotation, Visualization, and Integrated Discovery (DAVID) was then used to address the Gene Ontology Biological Process of significantly altered proteins between sources.

### Protein profile array

Secretory profiles were analyzed using a membrane-based antibody array (Human Neuro Discovery Array C1, RayBiotech, USA) according to the manufacturer’s instructions. Non-concentrated conditioned medium was incubated overnight at 4 °C, and bound proteins were detected via biotinylated antibody cocktail, streptavidin–HRP, and chemiluminescence (Sapphire Biomolecular Imager). Spot intensities were quantified using AzureSpot software (Azure Biosystems, USA).

### Animals & experimental design

Ten-weeks old Wistar-Han male rats (Charles River, Spain), weighing 305-414 g at the beginning of the experiment, were used, according to the Portuguese national authority for animal research, Direção Geral de Alimentação e Veterinária (ID: DGAV28421) and Ethical Subcommittee in Life and Health Sciences (SECVS; ID: SECVS 142/2016, University of Minho). Animals were housed in pairs, in a temperature and humidity-controlled room, maintained on 12 h light/dark cycles, with access to food and water *ad libitum*. Animals were handled for 1 week prior to experiments (Fig.2A). A total of 75 animals were used, divided into four groups as described in **Table 1**.

**Table 1.** Distribution and Description of Experimental Groups.

| Group | Lesion | Treatment | Number of animals (n) |
| --- | --- | --- | --- |
| Sham | None (0.2 mg/ml ascorbic acid) | Saline solution | 14 |
| 6-OHDA [Vehicle] | 6-OHDA [8µg] in the MFB | Neurobasal A medium | 19 |
| 6-OHDA [BM-MSCs CM] |  | BM-MSCs secretome | 21 |
| 6-OHDA [BM-MSCs<br>CM] |  | iMSCs secretome | 21 |

### Stereotaxic surgeries

Rats were anesthetized by intraperitoneal injection of ketamine (75 mg/kg) plus medetomidine (0.5 mg/kg) and placed on a stereotaxic apparatus (Stoelting, USA). The skull was exposed, and a burr hole was drilled. All surgeries were performed with a Hamilton syringe (30-gauge; Hamilton, Switzerland) at a flow rate of 0.5 μl /min. The lesioned group (n=62) was injected with a total of 8 μg of 6-OHDA (2 μl of 6-OHDA dissolved in saline containing 0.2 mg/ml ascorbic acid; Sigma, USA) in the right medial forebrain bundle (MFB) at the following coordinates related to Bregma: AP =4 .4 mm, ML = -1.0 mm, DV = -7.8 mm). The sham group (n=16) received an equal volume of vehicle solutions, at the same stereotaxic coordinates. The syringe was left in place for 2 minutes after injection and then removed slowly to optimize toxin diffusion and to avoid any backflow.

Treatment administration surgeries were performed as described above. The animals were unilaterally injected in the SNpc (4 μl, coordinates related to Bregma: AP = -5.3 mm, ML = 1.8 mm, DV = -7.4 mm) and into four different sites of the striatum (2 μl/site, coordinates related to Bregma: AP = -1.3 mm, ML = 4.7 mm, DV = -4.5 mm and -4.0 mm; AP = -0.4 mm, ML = 4.3 mm, DV = -4.5 mm and 4.0 mm; AP = 0.4 mm, ML = 3.1 mm, DV = -4.5 mm and -4.0 mm; AP = 1.3 mm, ML = 2.7 mm; DV = -4.5 mm and -4.0 mm). Postoperative analgesia consisted of buprenorphine (0.05 mg/kg, Bupaq, Richter Pharma, Wels, Austria).

### Behavioral analysis

#### Rotarod

Motor coordination and balance were assessed using a rotarod (TSE Systems, USA). Animals underwent 3 days of training on an accelerating protocol (4–40 rpm, 5 min), followed by a test day recording latency to fall. Trials were separated by ≥20 min rest.

#### Staircase

Fine motor control was evaluated using a staircase apparatus (Campden Instruments, USA) with food pellets placed on each step. Animals were familiarized for 10 min the day before testing and then allowed 15 min to retrieve pellets. Pellet consumption was recorded. Animals were food-restricted to 90% of free-feeding body weight to enhance motivation.

#### Apomorphine-induced rotations

Unilateral dopaminergic denervation was assessed by injecting apomorphine (0.5 mg/kg in saline containing 0.2% ascorbic acid) and recording net contralateral rotations over 30 min using an automated rotameter (MED-RSS, Med Associates, USA).

#### Sucrose consumption test

The sucrose consumption test was used to assess anhedonia. Pre-weighed drinking bottles containing water or 2% (m/v) sucrose solution were presented for 1 h to the animals individually placed on home cages. Before each test, rats were food and water- deprived for 12 h. Sucrose consumption test was performed during the nocturnal activity period of animals. Sucrose preference was calculated as previously described [76].

#### Sweet-drive test

Anhedonia was assessed using the Sweet Drive Test [77], a black acrylic arena divided into four chambers by transparent perforated walls. Rats were first placed in a pre- chamber connected via an automatic trapdoor to a middle chamber (MC). After entering the MC, the trapdoor closed, and animals could freely explore for 10 min between a right chamber containing 20 pre-weighed Cheerios® pellets (68% carbohydrates, 3800 kcal/kg) and a left chamber with 20 pre-weighed standard pellets (Mucedola 4RF21- GLP; 53.3% carbohydrates, 3700 kcal/kg). Sweet pellet preference (%) = [sweet pellet intake (g) / total intake (g)] × 100. Tests were performed after 10 h food deprivation, starting at 8:30 pm under red light.

Ultrasonic vocalizations (USVs, 50 kHz) were recorded using Avisoft CM16/CMPA microphones (10–200 kHz) positioned 20 cm above the floor and connected to an Avisoft UltrasoundGate 416H. Recordings were made with Avisoft-Recorder (v5.1.04; 250 kHz sampling rate, 16-bit format), and 50 kHz calls were quantified via automated analysis.

#### Elevated plus maze

The Elevated plus maze test was performed to address anxious-like behavior. It was carried out under a bright white light in an apparatus consisting of a black polypropylene plus-shaped platform elevated to 72.4 cm above the floor with two open arms (50.8 cm 10.2 cm) and two closed arms (50.8 cm 10.2 cm 40.6 cm; MedAssociates Inc., USA). Animals were placed individually in the center of the platform and the number of entries into each arm and the time spent therein were recorded during 5 min and analyzed using EthoVision XT (Noldus Information Technology, USA).

#### Novelty-suppressed feeding test

The test apparatus consisted of an open-field arena (MedAssociates Inc, USA) containing a single food pellet at the center. After a 18h food-deprivation period, animals were placed in the arena and the time the animal took to reach the pellet was measured and expressed as the latency to feed. After that, the animals were individually returned to their home cage, where they were allowed to eat a pre-weighed food pellet for 10 min to provide a measure of their appetite drive.

#### Tissue processing

At the end of behavioral testing, animals were euthanized with pentobarbital (Eutasil; 60 mg/kg, i.p.; Ceva Saúde Animal, Portugal). For histology (n = 6–8/group), rats were transcardially perfused with 0.9% saline followed by 4% PFA in PBS. Brains were post- fixed in 4% PFA for 48 h at RT, cryoprotected in 30% sucrose with 0.01% sodium azide at 4 °C, and sectioned coronally at 50 µm using a vibratome (VT1000S, Leica). For biochemical analyses (n = 10/group), brains were snap-frozen in liquid nitrogen, macro dissected using anatomical landmarks (Paxinos & Watson, 2005), and stored at −80 °C.

#### Histological analysis

Coronal sections containing dorsal striatum and NAc (5–6 series) and SNc and VTA (3– 4 series) were processed for free-floating TH immunohistochemistry. Sections were treated with 3% hydrogen peroxide (20 min), permeabilized in 0.1% PBS-T (10 min), and blocked in 10% NBCS/PBS (1 h). They were incubated overnight at 4 °C with rabbit anti-TH (1:1000; Merck Millipore), followed by biotinylated secondary antibody (goat anti-polyvalent; TP-125-BN, ThermoFisher Scientific) and streptavidin– peroxidase (TP-125-HR, ThermoFisher Scientific) for 30 min each at RT. Immunoreactivity was visualized with DAB (25 mg in 50 ml Tris-HCl 0.05 M, pH 7.6, plus 12.5 µl H[O[), sections were mounted, air-dried (24 h), counterstained with thionine, and coverslipped with Entellan™ (Merck Millipore).

TH+ fibers in the dorsal striatum and NAc were imaged under brightfield (SZX16, Olympus) and optical density was quantified in ImageJ (v1.48, NIH) relative to the non- lesioned side. TH+ cells in SNc and VTA were visualized (BX51, Olympus) and counted using Visiomorph (v2.12.3.0, Visiopharm) after delineating regional boundaries.

#### Neurochemical analysis

Monoaminergic neurotransmitters and metabolites in the striatum were quantified by HPLC with electrochemical detection (HPLC/EC; Gilson, Middleton, WI, USA) using a Supelcosil LC-18 column (3 mm; flow rate 1.0 ml/min). Left and right dorsal striata were weighed, homogenized in 0.2 N perchloric acid, sonicated on ice (5 min), and centrifuged at 5000 × g. Supernatants were filtered (Spin-X HPLC column, Costar) and 150 µl injected into the HPLC system. The mobile phase consisted of 0.7 M potassium phosphate buffer (pH 3.0) with 10% methanol, 1-heptanesulfonic acid (222 mg/L), and Na-EDTA (40 mg/L). Neurotransmitter concentrations were determined from daily standard curves and normalized to the amount of tissue from which they were extracted.

### SNc proteomic analysis

#### Sample preparation

For preparation of total proteome of the SNc, left and right SNc tissue obtained from microdissection were selected (three animals from each group). Brain tissues were lysed in 50 mM Trizma® hydrochloride solution pH 8 (Sigma) as described in [78]. Protein quantification using Pierce™ 660nm Protein Assay Reagent (Thermo Scientific™), 50 µg of total protein per sample (3 replicates per condition) was used for SWATH-MS. Four pooled samples were also created for protein identification/library creation by combining 15 µg of each replicate. All samples were spiked with equal amounts of internal standard (2 µg of MBP-GFP) prior protein digestion [69]. Samples were then digested as previously described [70].

#### Protein quantification by Sequential Windowed Acquisition of All Theoretical Mass Spectra (SWATH-MS)

Samples were analyzed on a NanoLC™ 425 system coupled to a TripleTOF™ 6600 mass spectrometer (Sciex®) in IDA and SWATH-MS modes, as for secretome analysis. For IDA, full MS scans (m/z 350–2250, 250 ms) were followed by up to 100 MS/MS scans (m/z 100–1500) per 3.295 s cycle, with precursor-dependent accumulation times (≥30 ms for >2000 intensity). Ions (+1 to +5, ≥10 cps) were fragmented once before 15 s dynamic exclusion. Rolling collision energy with CES 5 was applied. For SWATH, 168 overlapping windows (m/z 350–1250) were acquired with a 50 ms survey scan and 19 ms MS/MS scans, yielding 3.291 s cycles. Collision energy was set for +2 ions with variable CES.

For SWATH experiments, the mass spectrometer was operated in a looped product ion mode [71] and specifically tuned to a set of 168 overlapping windows, covering the precursor mass range of 350–1,250 m/z. A 50 ms survey scan (350–1,250 m/z) was acquired at the beginning of each cycle, and SWATH-MS/MS spectra were collected from 100 to 1,500 m/z for 19 ms resulting in a cycle time of 3.291 s. The collision energy for each window was determined according to the calculation for a charge +2 ion centered upon the window with variable CES, according to the window.

Peptide identification/library generation used ProteinPilot™ v5.1 (ABSciex®) against SwissProt *Rattus norvegicus* (Feb 2022) plus MBP-GFP sequences, with acrylamide- alkylated cysteines (fixed) and trypsin digestion. Positive identifications required ≤5% local FDR by target–decoy [72, 73].

Quantification was performed with the SWATH™ plug-in for PeakView™ (v2.0.01, ABSciex®) [74]. p to 15 peptides (≤5 fragments each) per protein were quantified if ≥1 confidence peptide (FDR <0.01) was present in ≥3 biological replicates per condition, with ≥3 transitions. XICs were extracted (4 min, 100 ppm). Protein levels were estimated by summing transitions across all qualifying peptides and normalized to total replicate intensity.

#### Bioinformatic analysis

Python was used to visualize patterns of protein abundance and perform differential protein analysis. Intensity values were log[-transformed and standardized per protein to z-scores (mean = 0, s.d. = 1). Hierarchical clustering (Euclidean distance, average linkage) was applied to both proteins and samples to explore overall expression patterns. To summarize global variation, principal component analysis (PCA) was performed on the transposed z-score matrix (samples × proteins) using scikit-learn. The first two principal components were plotted, with samples colored according to their experimental group.

Pairwise differential abundance testing between experimental groups (Sham, Vehicle, BM-MSCs, iMSCs) was conducted using two-sample Student’s t-tests assuming equal variance. Both raw p-values and Benjamini–Hochberg false discovery rate (FDR)– adjusted values were calculated. Significance was defined as p < 0.05 (or FDR < 0.05, where indicated) combined with an absolute log[fold-change cutoff of ≥ 0.5 (∼1.4- fold). Volcano plots were generated to visualize the relationship between fold-change and statistical significance. Data points were classified as significantly upregulated (positive log[ fold-change, p < 0.05), significantly downregulated (negative log[ fold- change, p < 0.05), or non-significant.

Lists of significantly altered proteins identified were uploaded into Ingenuity Pathway Analysis (IPA, QIAGEN Inc.) for functional annotation, namely, to identify enriched canonical pathways. Core analyses were performed using default parameters with species set to *Rattus norvegicu*s. Enrichment significance was determined using Fisher’s exact test (right tailed).

Finally, the list of proteins significantly altered in iMSCs secretome-treated animals versus non-treated animals was uploaded in STRING database (https://string-db.org) version 12.0. Protein–protein interaction was filtered at a medium to high confidence threshold (≥0.4) and further organized into three clusters, through DBSCAN clustering method (epsilon parameter = 3). Functional enrichment for KEGG pathways and Gene Ontology (GO) cellular components were computed using STRING’s built-in overrepresentation tests, applying a false discovery rate (FDR) < 0.05. The resulting interaction networks and pathway maps were exported for visualization and interpretation.

### Statistical analysis

Statistical analyses assumed a 95% confidence interval. Normality was assessed using histograms, skewness/kurtosis, and the Shapiro–Wilk test. Mean differences were evaluated using one-way ANOVA (single independent variable) or mixed-design ANOVA (one independent and one repeated-measures variable), with Tukey’s post hoc test for pairwise comparisons. Analyses were performed in GraphPad Prism 8.0.1, JASP 0.17.1, and Python.

## Abbreviations

6-OHDA: 6-hydroxydopamine
AMPK: AMP-activated protein kinase
asyn: α-synuclein
BIP / HSPA5: Endoplasmic reticulum chaperone BiP
BM-MSCs: Bone Marrow Mesenchymal Stem Cells
CALR: Calreticulin
CAP1: Adenylyl cyclase-associated protein 1
CFL1: Cofilin-1
DA: Dopamine
DCE2: Glutamate decarboxylase 2
ECM: Extracellular matrix
EEF2: Eukaryotic Elongation Factor 2
eIF4: Eukaryotic Initiation Factor 4
EIF4A1: Eukaryotic initiation factor 4A-I
EIF5A: Eukaryotic translation initiation factor 5A
ENO1: Enolase 1
EPM: Elevated Plus Maze
FKBP10: Peptidyl-prolyl cis-trans isomerase
FN1: Fibronectin
GABR1: Gamma-aminobutyric acid type B receptor subunit 1
GAPDH: Glyceraldehyde-3-phosphate dehydrogenase
Gnb1: G Protein Subunit Beta 1
Gnb4: G Protein Subunit Beta 4
Gpx3: Glutathione peroxidase 3
GRIA2: Glutamate receptor 2
Gstm2: Glutathione S-transferase Yb-3
Gstm3: Glutathione S-transferase Mu 5
GSTP1: Glutathione S-transferase P
H2AC18: Histone H2A type 2-A
H3C13: Histone H3.2
H4C6: H4 clustered histone 6
HB-EGF: Heparin-Binding Epidermal Growth Factor-Like Growth Factor
HSP90AB1: Heat shock protein 90 alpha family class B member 1
HSP90B1: Heat Shock Protein 90 Beta Family Member 1
HSPA5: Heat shock protein family A member 5
HSPA8: Heat shock protein family A member 8
IFNγ: Interferon gamma
IGF-1: Insulin-Like Growth Factor 1
IGFBP7 / IBP7: Insulin-like growth factor-binding protein 7
IL-10: Interleukin 10
IL-1α: Interleukin 1 alpha
IL-1β: Interleukin 1 beta
IL-6: Interleukin-6
IL-8: Interleukin-8
iPSCs: induced Pluripotent Stem Cells
Keap1/Nrf2/ARE: 1/Nuclear factor-erythroid-2-related factor 2/Antioxidant response element
LDHA: Lactate dehydrogenase A
LMNA: Prelamin-A/C
MFB: Medial Forebrain bundle
MIP-1α: Macrophage Inflammatory Protein-1 alpha
MMP2: Matrix metallopeptidase 2
MMP3: Matrix metallopeptidase 3
MOBP: Myelin-associated oligodendrocyte basic protein ()
MPTP: 1-methyl-4-phenyl-1,2,3,6-tetrahydropyridine
MSCs: Mesenchymal Stem Cells
mTOR: Mechanistic Target of Rapamycin
MYH9: Myosin-9
NAc: *nucleus accumbens*
NSF: Novelty Suppressed Feeding
P4HB: Prolyl 4-hydroxylase subunit beta
p70S6K: 70 kDa Ribosomal Protein S6 Kinase
PD: Parkinson’s Disease
PDIA1: Protein disulfide-isomerase A1
PDIA3: Protein disulfide-isomerase A3
PFN1: Profilin-1
PFN2: Profilin-2
PKA: Protein Kinase A
PKM: Pyruvate kinase
PPIB: Peptidyl-prolyl cis-trans isomerase B
PRDX1: Peroxidoxin-1
Prkacb: Protein Kinase cAMP-Activated Catalytic Subunit Beta
PSMA6: Proteasome subunit alpha type-6
PSMB11: proteasome subunit beta type-11
PSMB2: proteasome subunit beta type-2
PSMB4: proteasome subunit beta type-4
PSMB6: Proteasome subunit beta type-6
PSME1: proteasome activator complex subunit 1
PSMG2: proteasome assembly chaperone 2
PSMG4: proteasome assembly chaperone 4
Rap1b: Ras-Related Protein Rap-1b
Rhoa: Ras Homolog Family Member A
ROS: Reactive oxygen species
RPL14: Large ribosomal subunit protein eL14
RPS11: Ribosomal protein S11
RPS18: Ribosomal protein S18
RPS23: Ribosomal protein S23
RPS9: Ribosomal protein S9
S100B: Calcium Binding Protein B
SDC1: Syndecan-1
SNc: *Substancia nigra pars compacta*
TGF-β: Growth Factor Beta
TGFα: Transforming Growth Factor alpha
TH: Tyrosine hydroxylase
THIO: Thioredoxin
TIMP1: Metalloproteinase inhibitor 1
TPI1: Triosephosphate isomerase
TPM1: Tropomyosin alpha-1 chain
TUB1A: alpha-tubulin-1A
TUBA1C: Tubulin alpha-1C chain
TUBB: Tubulin beta chain
TUBB2B: Tubulin beta-2B chain
TUBB4B: Tubulin beta-4B chain
TUBB6: Tubulin beta-6 chain
UBA52: Ubiquitin-ribosomal protein eL40 fusion protein
ULK1: Unc-51 **l**ike **a**utophagy **a**ctivating **k**inase 1
USVs: Ultrasonic vocalizations
VEGF-A: Vascular Endothelial Growth Factor A
VGLU2: Vesicular glutamate transporter 2
VIM / VIME: Vimentin
VTA: Ventral tegmental area
YWHAZ / 14-3-3ζ / 1433Z: 14-3-3 protein zeta/delta

## Data availability statement

(All published manuscripts reporting original research must include a data availability statement. The data availability statement must make the conditions of access to the minimum dataset that are necessary to interpret, verify and extend the research in the article transparent to readers.

## Acknowledgments

This work was funded by “la Caixa” Foundation and Fundação para a Ciência e a Tecnologia (FCT), through Iniciativa Ibérica de Investigación y Innovación Biomedica, under the agreement LCF/PR/HP20/52300001; by National funds, through FCT – project UIDB/50026/2020, UIDP/50026/2020 and 2023.13115.PEX.; by the project NORTE-01-0145-FEDER-000039, supported by Norte Portugal Regional Operational Programme (NORTE 2020), under the PORTUGAL 2020 Partnership Agreement, through the European Regional Development Fund (ERDF); Additional support was provided by the FCT through a PhD Fellowship awarded to FFA (UI/BD/154989/2023).

## Author contributions

A.M. and A.J.S. conceived the study. A.M., F.F.A., S.B.A., and S.I.A. designed and performed methodology. A.M., F.F.A., B.S.M., and A.J.S. validated the data. A.M., F.F.A., and S.I.A. performed formal analysis, and A.M., F.F.A., S.B.A., B.M.P, C.S.C., A.V.D., conducted the investigation. B.M., C.S.C, provided resources. S.I.A. curated the data. A.M. and F.F.A. wrote the original draft. A.M., F.F.A., B.S.M., and A.J.S. reviewed and edited the manuscript. A.M., F.F.A., and S.I.A. prepared the visualizations. B.S.M. and A.J.S. supervised the project. Project administration and funding acquisition were led by A.J.S.

## References

1. Beitz, J.M., Parkinson’s disease: a review. Front Biosci (Schol Ed), 2014. 6(1): p. 65–74.

2. Pires, A.O., et al., Old and new challenges in Parkinson’s disease therapeutics. Prog Neurobiol, 2017. 156: p. 69–89.

3. Athauda, D. and T. Foltynie, The ongoing pursuit of neuroprotective therapies in Parkinson disease. Nat Rev Neurol, 2015. 11(1): p. 25–40.

4. Volkman, R. and D. Offen, Concise Review: Mesenchymal Stem Cells in Neurodegenerative Diseases. Stem Cells, 2017. 35(8): p. 1867–1880.

5. Marques, C.R., et al., Secretome of bone marrow mesenchymal stromal cells cultured in a dynamic system induces neuroprotection and modulates microglial responsiveness in an α-synuclein overexpression rat model. Cytotherapy, 2024. 26(7): p. 700–713.

6. Marques, C.R., et al., Mesenchymal stem cell secretome protects against alpha- synuclein-induced neurodegeneration in a Caenorhabditis elegans model of Parkinson’s disease. Cytotherapy, 2021. 23(10): p. 894–901.

7. Teixeira, F.G., et al., Impact of the Secretome of Human Mesenchymal Stem Cells on Brain Structure and Animal Behavior in a Rat Model of Parkinson’s Disease. Stem Cells Transl Med, 2017. 6(2): p. 634–646.

8. Mendes-Pinheiro, B., et al., Treating Parkinson’s Disease with Human Bone Marrow Mesenchymal Stem Cell Secretome: A Translational Investigation Using Human Brain Organoids and Different Routes of In Vivo Administration. Cells, 2023. 12(21).

9. Mendes-Pinheiro, B., et al., Bone Marrow Mesenchymal Stem Cells’ Secretome Exerts Neuroprotective Effects in a Parkinson’s Disease Rat Model. Front Bioeng Biotechnol, 2019. 7: p. 294.

10. Marote, A., et al., Cellular Aging Secretes: a Comparison of Bone-Marrow- Derived and Induced Mesenchymal Stem Cells and Their Secretome Over Long- Term Culture. Stem Cell Rev Rep, 2023. 19(1): p. 248–263.

11. Pinho, A.G., et al., Immunomodulatory and regenerative effects of the full and fractioned adipose tissue derived stem cells secretome in spinal cord injury. Exp Neurol, 2022. 351: p. 113989.

12. Fan, X.L., et al., Mechanisms underlying the protective effects of mesenchymal stem cell-based therapy. Cell Mol Life Sci, 2020. 77(14): p. 2771–2794.

13. Fan, X.-L., et al., Mechanisms underlying the protective effects of mesenchymal stem cell-based therapy. Cell Mol Life Sci, 2020: p. 10.1007/s00018-020-03454-6.

14. Fitzsimmons, R.E.B., et al., Mesenchymal Stromal/Stem Cells in Regenerative Medicine and Tissue Engineering. Stem Cells Int, 2018. 2018: p. 8031718–8031718.

15. Hynes, K., et al., Generation of functional mesenchymal stem cells from different induced pluripotent stem cell lines. Stem Cells Dev, 2014. 23(10): p. 1084–96.

16. Sabapathy, V. and S. Kumar, hiPSC-derived iMSCs: NextGen MSCs as an advanced therapeutically active cell resource for regenerative medicine. Journal of Cellular and Molecular Medicine, 2016. 21(10): p. 128–39.

17. Zhou, X., et al., The application potential of iMSCs and iMSC-EVs in diseases. Front Bioeng Biotechnol, 2024. 12: p. 1434465.

18. Ren, H., et al., The therapeutic effects of induced pluripotent stem cell-derived mesenchymal stem cells on Parkinson’s disease. IUBMB Life, 2025. 77(1): p. e2936.

19. Pires, A.O., et al., Unveiling the Differences of Secretome of Human Bone Marrow Mesenchymal Stem Cells, Adipose Tissue-Derived Stem Cells, and Human Umbilical Cord Perivascular Cells: A Proteomic Analysis. Stem Cells Dev, 2016. 25(14): p. 1073–83.

20. Assunção-Silva, R.C., et al., Exploiting the impact of the secretome of MSCs isolated from different tissue sources on neuronal differentiation and axonal growth. Biochimie, 2018. 155: p. 83–91.

21. Rike, W.A. and S. Stern, Proteins and Transcriptional Dysregulation of the Brain Extracellular Matrix in Parkinson’s Disease: A Systematic Review. Int J Mol Sci, 2023. 24(8).

22. Hu, S., et al., Molecular chaperones and Parkinson’s disease. Neurobiol Dis, 2021. 160: p. 105527.

23. Hartl, F.U., A. Bracher, and M. Hayer-Hartl, Molecular chaperones in protein folding and proteostasis. Nature, 2011. 475(7356): p. 324–32.

24. Carvalho, M.M., et al., Behavioral characterization of the 6-hydroxidopamine model of Parkinson’s disease and pharmacological rescuing of non-motor deficits. Mol Neurodegener, 2013. 8: p. 14.

25. Burke, R.E., et al., An assessment of the validity of densitometric measures of striatal tyrosine hydroxylase-positive fibers: relationship to apomorphine- induced rotations in 6-hydroxydopamine lesioned rats. J Neurosci Methods, 1990. 35(1): p. 63–73.

26. Magnard, R., et al., What can rodent models tell us about apathy and associated neuropsychiatric symptoms in Parkinson’s disease? Transl Psychiatry, 2016. 6(3): p. e753.

27. Mateus-Pinheiro, A., et al., The Sweet Drive Test: refining phenotypic characterization of anhedonic behavior in rodents. Front Behav Neurosci, 2014. 8: p. 74.

28. Portfors, C.V., Types and functions of ultrasonic vocalizations in laboratory rats and mice. J Am Assoc Lab Anim Sci, 2007. 46(1): p. 28–34.

29. Wei, L., et al., Abnormal ventral tegmental area-anterior cingulate cortex connectivity in Parkinson’s disease with depression. Behav Brain Res, 2018. 347: p. 132–139.

30. Alberico, S.L., M.D. Cassell, and N.S. Narayanan, The Vulnerable Ventral Tegmental Area in Parkinson’s Disease. Basal Ganglia, 2015. 5(2-3): p. 51–55.

31. Zhao, J., et al., Multidimensional mechanisms of anxiety and depression in Parkinson’s disease: Integrating neuroimaging, neurocircuits, and molecular pathways. Pharmacol Res, 2025. 215: p. 107717.

32. Li, Y., P. Maher, and D. Schubert, Requirement for cGMP in nerve cell death caused by glutathione depletion. J Cell Biol, 1997. 139(5): p. 1317–24.

33. Moradi Vastegani, S., et al., Mitochondrial Dysfunction and Parkinson’s Disease: Pathogenesis and Therapeutic Strategies. Neurochem Res, 2023. 48(8): p. 2285–2308.

34. Trist, B.G., D.J. Hare, and K.L. Double, Oxidative stress in the aging substantia nigra and the etiology of Parkinson’s disease. Aging Cell, 2019. 18(6): p. e13031.

35. Bjørklund, G., et al., The glutathione system in Parkinson’s disease and its progression. Neurosci Biobehav Rev, 2021. 120: p. 470–478.

36. Alharbi, B., et al., Role of GABA pathway in motor and non-motor symptoms in Parkinson’s disease: a bidirectional circuit. Eur J Med Res, 2024. 29(1): p. 205.

37. Fortin, G.M., et al., Glutamate corelease promotes growth and survival of midbrain dopamine neurons. J Neurosci, 2012. 32(48): p. 17477–91.

38. Choudhury, S.P., et al., Altered neural cell junctions and ion-channels leading to disrupted neuron communication in Parkinson’s disease. NPJ Parkinsons Dis, 2022. 8(1): p. 66.

39. Li, X., et al., Association of Myelin Disruption and Iron Accumulation on MRI With Parkinson’s Disease Severity. J Magn Reson Imaging, 2025.

40. Merimi, M., et al., The Therapeutic Potential of Mesenchymal Stromal Cells for Regenerative Medicine: Current Knowledge and Future Understandings. Front Cell Dev Biol, 2021. 9: p. 661532.

41. Wruck, W., et al., Human Induced Pluripotent Stem Cell-Derived Mesenchymal Stem Cells Acquire Rejuvenation and Reduced Heterogeneity. Front Cell Dev Biol, 2021. 9: p. 717772.

42. Beutgen, V.M., et al., Secretome Analysis Using Affinity Proteomics and Immunoassays: A Focus on Tumor Biology. Mol Cell Proteomics, 2024. 23(9): p. 100830.

43. Palamà, M.E.F., et al., Batch variability and anti-inflammatory effects of iPSC- derived mesenchymal stromal cell extracellular vesicles in osteoarthritis. Front Bioeng Biotechnol, 2025. 13: p. 1536843.

44. Chouaib, B., et al., Towards the Standardization of Mesenchymal Stem Cell Secretome-Derived Product Manufacturing for Tissue Regeneration. Int J Mol Sci, 2023. 24(16).

45. Marote, A., et al., MSCs-Derived Exosomes: Cell-Secreted Nanovesicles with Regenerative Potential. Front Pharmacol, 2016. 7: p. 231.

46. Vizoso, F.J., et al., Mesenchymal Stem Cell Secretome: Toward Cell-Free Therapeutic Strategies in Regenerative Medicine. Int J Mol Sci, 2017. 18(9).

47. Padhan, P., et al., Glutathione S-transferase: A keystone in Parkinson’s disease pathogenesis and therapy. Mol Cell Neurosci, 2025. 132: p. 103981.

48. Allen, M., et al., Glutathione S-transferase omega genes in Alzheimer and Parkinson disease risk, age-at-diagnosis and brain gene expression: an association study with mechanistic implications. Mol Neurodegener, 2012. 7: p. 13.

49. Alnasser, S.M., The role of glutathione S-transferases in human disease pathogenesis and their current inhibitors. Genes Dis, 2025. 12(4): p. 101482.

50. Lastres-Becker, I., et al., Repurposing the NRF2 Activator Dimethyl Fumarate as Therapy Against Synucleinopathy in Parkinson’s Disease. Antioxid Redox Signal, 2016. 25(2): p. 61–77.

51. Ho, H.K., et al., Nrf2 activation involves an oxidative-stress independent pathway in tetrafluoroethylcysteine-induced cytotoxicity. Toxicol Sci, 2005. 86(2): p. 354–64.

52. Quintanilla, M.E., et al., Intranasal mesenchymal stem cell secretome administration markedly inhibits alcohol and nicotine self-administration and blocks relapse-intake: mechanism and translational options. Stem Cell Res Ther, 2019. 10(1): p. 205.

53. Malagelada, C., et al., Rapamycin protects against neuron death in in vitro and in vivo models of Parkinson’s disease. J Neurosci, 2010. 30(3): p. 1166–75.

54. Zhou, Q., et al., Sulforaphane protects against rotenone-induced neurotoxicity in vivo: Involvement of the mTOR, Nrf2, and autophagy pathways. Sci Rep, 2016. 6: p. 32206.

55. Lan, A.P., et al., mTOR Signaling in Parkinson’s Disease. Neuromolecular Med, 2017. 19(1): p. 1–10.

56. Panwar, V., et al., Multifaceted role of mTOR (mammalian target of rapamycin) signaling pathway in human health and disease. Signal Transduct Target Ther, 2023. 8(1): p. 375.

57. Zhu, Z., et al., Balancing mTOR Signaling and Autophagy in the Treatment of Parkinson’s Disease. Int J Mol Sci, 2019. 20(3).

58. Behl, T., et al., Distinctive Evidence Involved in the Role of Endocannabinoid Signalling in Parkinson’s Disease: A Perspective on Associated Therapeutic Interventions. Int J Mol Sci, 2020. 21(17).

59. Stampanoni Bassi, M., et al., Cannabinoids in Parkinson’s Disease. Cannabis Cannabinoid Res, 2017. 2(1): p. 21–29.

60. Basavarajappa, B.S., et al., Endocannabinoid system in neurodegenerative disorders. J Neurochem, 2017. 142(5): p. 624–648.

61. Lowe, H., et al., The Endocannabinoid System: A Potential Target for the Treatment of Various Diseases. Int J Mol Sci, 2021. 22(17).

62. Walsh, S., et al., Loss of cannabinoid CB1 receptor expression in the 6- hydroxydopamine-induced nigrostriatal terminal lesion model of Parkinson’s disease in the rat. Brain Res Bull, 2010. 81(6): p. 543–8.

63. Sedaghat, K., A.L. Gundlach, and D.I. Finkelstein, Analysis of morphological and neurochemical changes in subthalamic nucleus neurons in response to a unilateral 6-OHDA lesion of the substantia nigra in adult rats. IBRO Neurosci Rep, 2021. 10: p. 96–103.

64. Dzamko, N., Cytokine activity in Parkinson’s disease. Neuronal Signal, 2023. 7(4): p. NS20220063.

65. Zhang, R., et al., Anti-inflammatory and immunomodulatory mechanisms of mesenchymal stem cell transplantation in experimental traumatic brain injury. J Neuroinflammation, 2013. 10: p. 106.

66. Mauri, M., et al., Mesenchymal stem cells enhance GABAergic transmission in co-cultured hippocampal neurons. Mol Cell Neurosci, 2012. 49(4): p. 395–405.

67. Salazar Campos, J.M., L.F. Burbulla, and S. Jäkel, Are oligodendrocytes bystanders or drivers of Parkinson’s disease pathology? PLoS Biol, 2025. 23(1): p. e3002977.

68. Vilaça-Faria, H., et al., Fractionating stem cells secretome for Parkinson’s disease modeling: Is it the whole better than the sum of its parts? Biochimie, 2021. 189: p. 87–98.

69. Anjo, S.I., et al., Use of recombinant proteins as a simple and robust normalization method for untargeted proteomics screening: exhaustive performance assessment. Talanta, 2019. 205: p. 120163.

70. Anjo, S.I., C. Santa, and B. Manadas, Short GeLC-SWATH: A fast and reliable quantitative approach for proteomic screenings. PROTEOMICS, 2015. 15(4): p. 757–762.

71. Gillet, L.C., et al., Targeted Data Extraction of the MS/MS Spectra Generated by Data-independent Acquisition: A New Concept for Consistent and Accurate Proteome Analysis*. Molecular & Cellular Proteomics, 2012. 11(6): p. O111.016717.

72. Sennels, L., J.-C. Bukowski-Wills, and J. Rappsilber, Improved results in proteomics by use of local and peptide-class specific false discovery rates. BMC Bioinformatics, 2009. 10: p. 179–179.

73. Tang, W.H., I.V. Shilov, and S.L. Seymour, Nonlinear fitting method for determining local false discovery rates from decoy database searches. J Proteome Res., 2008. 7(9): p. 3661–7. doi: 10.1021/pr070492f. Epub 2008 Aug 14.

74. Lambert, J.-P., et al., Mapping differential interactomes by affinity purification coupled with data-independent mass spectrometry acquisition. Nature methods, 2013. 10(12): p. 1239–1245.

75. Collins, B.C., et al., Quantifying protein interaction dynamics by SWATH mass spectrometry: application to the 14-3-3 system. Nat Methods., 2013. 10(12): p. 1246–53. doi: 10.1038/nmeth.2703. Epub 2013 Oct 27.

76. Bessa, J., et al., A trans-dimensional approach to the behavioral aspects of depression. Front Behav Neurosci, 2009. 3(1).

77. Mateus-Pinheiro, A., et al., The Sweet Drive Test: refining phenotypic characterization of anhedonic behavior in rodents. Front Behav Neurosci, 2014. 8: p. 74–74.

78. Anjo, S.I., et al., Neuroproteomics Using Short GeLC-SWATH: From the Evaluation of Proteome Changes to the Clarification of Protein Function, in Current Proteomic Approaches Applied to Brain Function, E. Santamaría and J. Fernández-Irigoyen, Editors. 2017, Springer New York: New York, NY. p. 107–138.

