## Supplementary for "iPSC-derived MSC secretome activates distinct neuroprotective pathways in a preclinical model of Parkinson’s disease"

### Supplementary Figure List

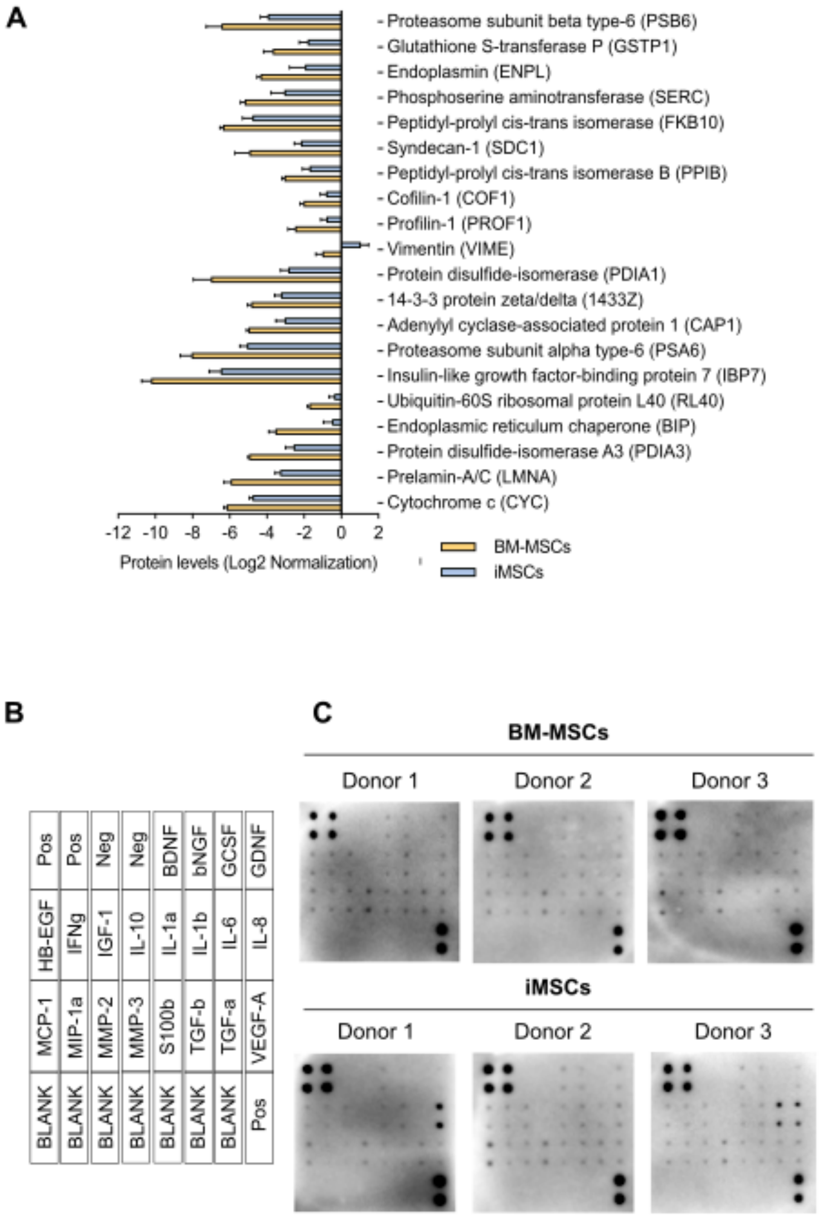


**Supplemenatary Figure 1.A.** Protein levels of significantly altered proteins between sources after Log2 Normalization; **B**. Human Neuro Discovery membrane array map identifying the position of each antibody on the membrane; **C**. Membrane arrays depicting the proteins detected on the conditioned medium of each donor from BM-MSCs and iMSCs.

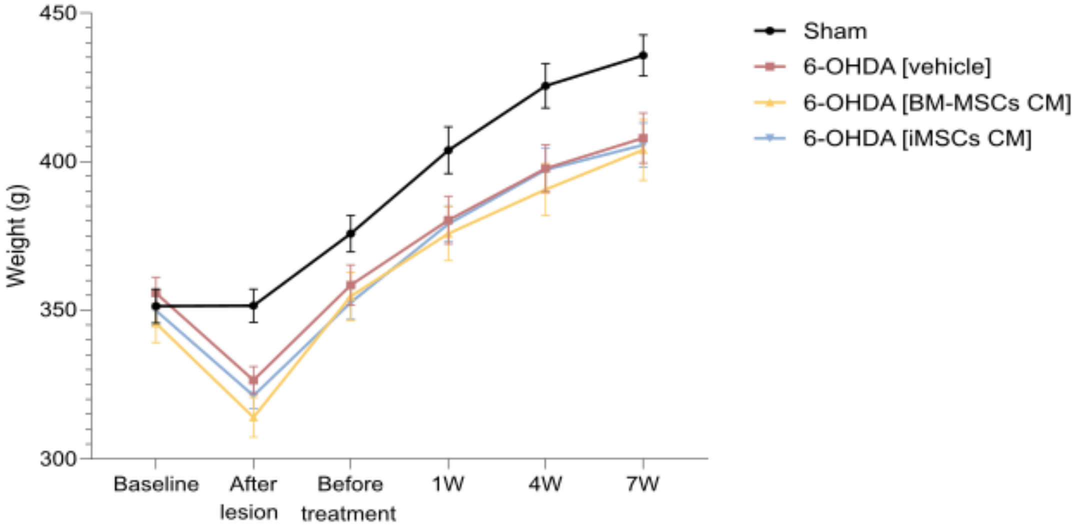


**Supplemenatary Figure 2.** Body weight variation starting from baseline, after lesion, before treatment and 1-, 4- and 7- weeks after treatment analyzed using a two-way ANOVA with time (row factor) and weight (column factor) and their interaction as sources of variation. Significant main effects were observed for the column factor ($F$(3, 62) = 2.815, $P$ = 0.0464) and the row factor ($F$(2.617, 162.2) = 604.3, $P$ < 0.001), as well as for the interaction between factors ($F$(15, 310) = 4.749, $P$ <0.0001).

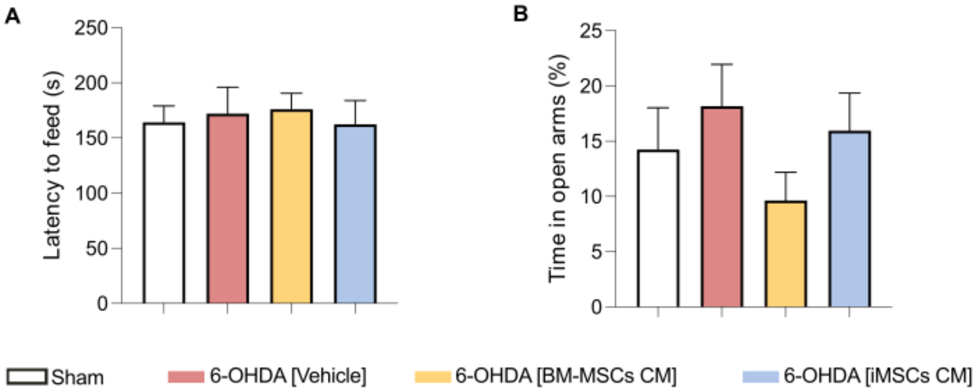


**Supplementary Figure 3**. **A**. Latency to feed as a measure of the novelty suppressed feeding test analyzed using a Kruskal–Wallis test, which revealed no significant differences among groups (Kruskal–Wallis statistic = 0.6568, $P$ = 0.8833; $n$ = 4 groups, 64 total values). **B**. time spent in open arms on the elevated plus maze test analyzed using Brown–Forsythe and Welch's ANOVA tests to account for unequal variances across treatments. Brown–Forsythe ($F(3, 48.45) = 1.103$, $P = 0.3570$) and Welch's ANOVA ($W(3, 28.17) = 1.387$, $P = 0.2673$).

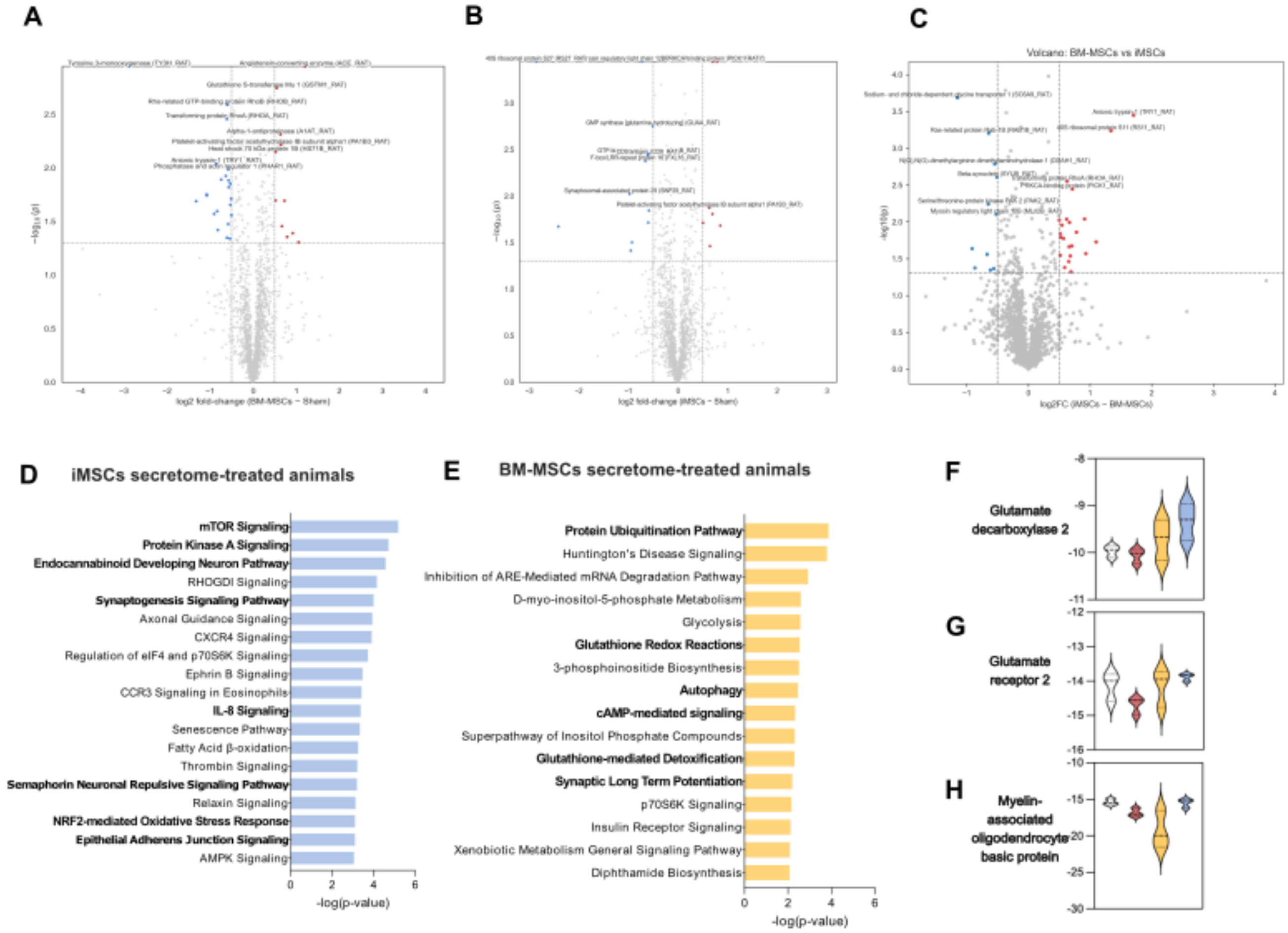


**Supplementary Figure 4. A.** Volcano plot displaying differential protein abundance between Vehicle and BM-MSCs secretome treated animals. Significantly altered proteins were defined as those with Benjamini–Hochberg FDR–adjusted p value < 0.05 in pairwise t-tests and an absolute log2 fold-change threshold of ≥ 0.5 (~1.4-fold) and are represented in the graph. **B**. Volcano plot displaying differential protein abundance between Sham and BM-MSCs secretome treated animals Significantly altered proteins were defined as those with Benjamini–Hochberg FDR–adjusted p value < 0.05 in pairwise t-tests and an absolute log2 fold-change threshold of ≥ 0.5 (~1.4-fold) and are represented in the graph. **C**. Volcano plot displaying differential protein abundance between iMSCs and BM-MSCs secretome treated animals. Significantly altered proteins were defined as those with Benjamini–Hochberg FDR–adjusted p value < 0.05 in pairwise t-tests and an absolute log2 fold-change threshold of ≥ 0.5 (~1.4-fold) and are represented in the graph. **D.** Over-representation analysis of cellular component terms using

ConsensusPathDB, highlighting enrichment in protein-folding-related components found in brain tissue of BM-MSCs treated rats. **E.** Over-representation analysis of cellular component terms using ConsensusPathDB, highlighting enrichment in protein-folding-related components found in brain tissue of BM-MSCs treated rats **F**. Log2 normalized values of Glutamate decarboxylase 2. **G**. Log2 normalized values of Glutamate receptor 2. **H**. Log2 normalized values of Myelin-associated oligodendrocyte basic protein.
